# Using the past to estimate sensory uncertainty

**DOI:** 10.1101/589275

**Authors:** Ulrik Beierholm, Tim Rohe, Ambra Ferrari, Oliver Stegle, Uta Noppeney

## Abstract

To form the most reliable percept of the environment, the brain needs to represent sensory uncertainty. Current theories of perceptual inference assume that the brain computes sensory uncertainty instantaneously and independently for each stimulus.

In a series of psychophysics experiments human observers localized auditory signals that were presented in synchrony with spatially disparate visual signals. Critically, the visual noise changed dynamically over time with or without intermittent jumps. Our results show that observers integrate audiovisual inputs weighted by sensory reliability estimates that combine information from past and current signals as predicted by an optimal Bayesian learner or approximate strategies of exponential discounting

Our results challenge classical models of perceptual inference where sensory uncertainty estimates depend only on the current stimulus. They demonstrate that the brain capitalizes on the temporal dynamics of the external world and estimates sensory uncertainty by combining past experiences with new incoming sensory signals.

---

Perception has been described as a process of statistical inference based on noisy sensory inputs^1^. Key to this perceptual inference is the representation of sensory uncertainty. Most prominently, in multisensory perception the most reliable or ‘Bayes-optimal’ percept is obtained by integrating sensory signals weighted by their reliability (i.e., precision or inverse of variance) with less weight assigned to less reliable signals. Indeed, accumulating evidence suggests that human observers are close to optimal in many perceptual tasks (though see ^1–4^) and weight signals according to their sensory reliabilities^5–9^.

An unresolved question is how human observers compute and represent their sensory uncertainty. Current theories and experimental approaches generally assume that observers access sensory uncertainty near-instantaneously and independently across briefly (≤200 ms) presented stimuli ^10,11^. At the neural level, theories of probabilistic population coding have suggested that sensory uncertainty may be represented instantaneously in the gain of the neuronal population response^12^. Yet, in our natural environment sensory noise often evolves at slow timescales. For instance, visual noise slowly varies when walking through a snow storm. Observers may capitalize on the temporal dynamics of the external world and use the past to inform current estimates of sensory uncertainty. In this alternative account, more reliable estimates of sensory uncertainty would be obtained by combining past estimates with current sensory inputs as predicted by Bayesian learning.

To arbitrate between these two critical hypotheses, we presented observers with audiovisual signals in synchrony but with a small spatial disparity in a sound localization task. Critically, the spatial standard deviation (STD) of the visual signal changed dynamically over time continuously (experiment 1-3) or discontinuously (i.e. with intermittent jumps; experiment 4). First, we investigated whether the influence of the visual signal location on observers’ perceived sound location depended on the noise only of the current visual signal or also of past visual signals. Second, using computational modeling and Bayesian model comparison, we formally assessed whether observers update their visual uncertainty estimates consistent with i. an instantaneous learner, ii. an optimal Bayesian learner or iii. an exponential learner.

## Results

In a series of four experiments, we presented participants with audiovisual signals in a spatial localization task where physical visual noise changed dynamically over time at 5 Hz either continuously (experiments 1-3) or discontinuously with intermittent jumps (experiment 4, Fig. 1). On each trial, participants located the sound that was presented in synchrony with a change in color of the visual cloud of dots. The spatial disparity between the auditory signal and the mean of the visual cloud was set to ± 5° visual angle. This small audiovisual disparity enabled an influence of the visual signal location on the perceived sound location as a function of visual noise^5,13^. As a result, observers’ visual uncertainty estimate could be quantified in terms of the relative weight of the auditory signal on the perceived sound location with a greater auditory weight indicating that observers estimated a greater visual uncertainty. Importantly, while the visual cloud’s mean (together with the corresponding sound position) was independently resampled on each trial from five possible locations, the visual cloud was re-displayed at a higher temporal rate of 5 Hz with the cloud’s horizontal standard deviation (STD) slowly varying over time.

**Figure 1.**
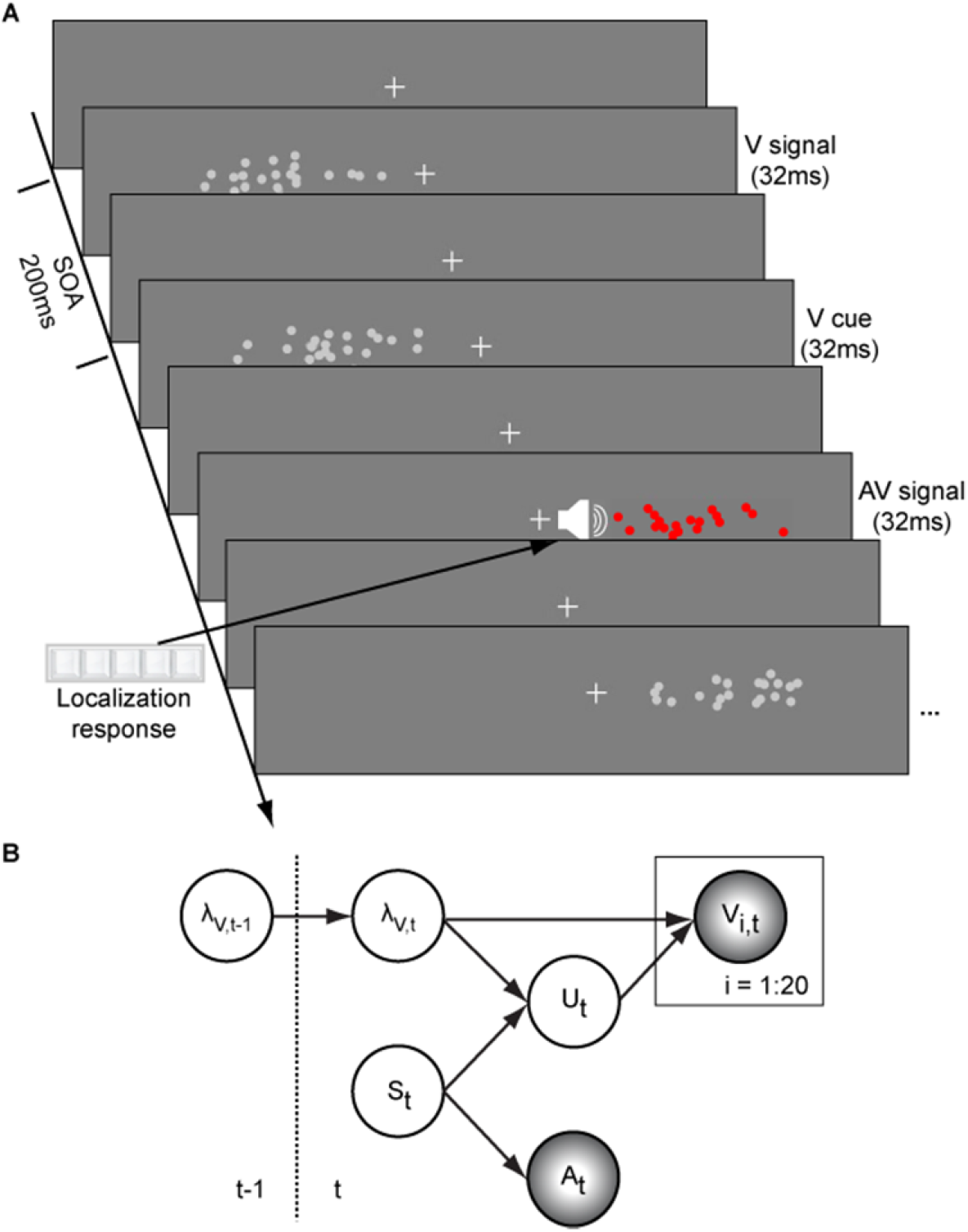
Audiovisual localization paradigm and Bayesian causal inference model for learning visual reliability. (**A**) Visual (V) signals (cloud of 20 bright dots) were presented at 5 Hz (i.e. every 200 ms). The cloud’s location mean was temporally independently resampled from five possible locations (−10°, −5°, 0°, 5°, 10°) at a SOA jittered between 1.4 and 2.8 s. In synchrony with the change in the cloud’s location, the dots changed their colour and a sound was presented (AV signal) which the participants localized using five response buttons. The location of the sound was sampled from the two possible locations adjacent to the visual cloud’s mean location (i.e. ± 5° AV spatial discrepancy). (**B**) The generative model for the Bayesian learner explicitly modelled the potential causal structures, i.e. whether visual (V_i_) signals and an auditory (A) signal are generated by one common audiovisual source S_t_, i.e. C = 1, or by two independent sources S_Vt_ and S_At_, i.e. C = 2 (n.b. only the model component for the common source case is shown to illustrate the temporal updating, for complete generative model, see Supplementary Fig. 1). Importantly, the reliability (i.e., 1/variance) of the visual signal at time t (λ_t_) depends on the reliability of the previous visual signal (λ_t−1_) for both model components (i.e. common source and independent source).

In the first three experiments, we used continuous sequences, where the visual cloud’s STD changed periodically according to a sinusoid (n = 25; period = 30 s), a random walk (RW1; n = 33; period = 120 s) or a smoothed random walk (RW2; n = 19; period = 30 s; Fig. 2). In an additional fourth experiment, we inserted abrupt increases or decreases into a sinusoidal evolution of the visual cloud’s STD (n =18, period = 30 s, Fig. 4). We will first describe the results for the three continuous sequences followed by the discontinuous sequence.

**Figure 2.**
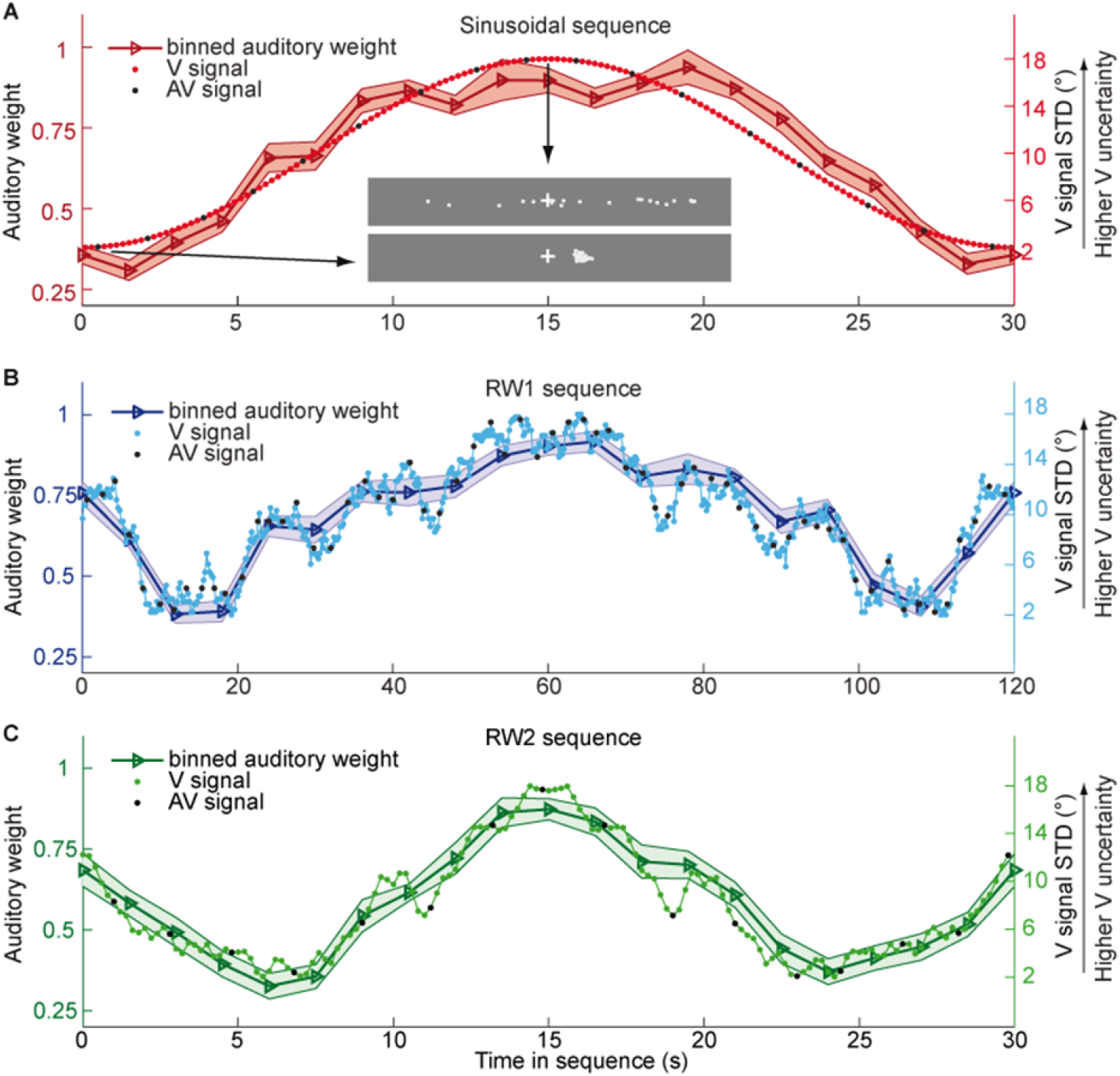
Time course of visual noise and relative auditory weights for continuous sequences of visual noise. Relative auditory weights (mean across participants ± SEM, left ordinate) and visual noise (i.e., STD of the cloud of dots, right ordinate) are displayed as a function of time. The relative auditory weight varies between one (i.e. pure auditory influence on the localization responses) and zero (i.e. pure visual influence). The STD of the visual cloud was manipulated as (**A**) a sinusoidal (period 30s, N = 25), (**B**) a random walk (RW1, period 120s, N = 33) and (**C**) a smoothed random walk (RW2, period 30s, N = 19). Please note that the period for RW1 sequence is 120 s, while the periods of the sinusoidal and RW2 is only 30 s. As a result, the overall dynamics as quantified by the power spectrum is faster for RW2 than RW1 (peak in frequency range [0 0.2] Hz: Sinusoid: 0.033 Hz, RW1: 0.025 Hz, RW2: 0.066 Hz). The sequence of visual clouds was presented at 5 Hz, while trials, i.e. audiovisual (AV) signals (colour change with sound presentation, black dots), were interspersed with a SOA jittered between 1.4 and 2.8 s. For illustration purposes, the cloud of dots for the lowest (i.e., V signal STD = 2°) and the highest (i.e., V signal STD = 18°) visual variance are shown in (**A**).

**Figure 3.**
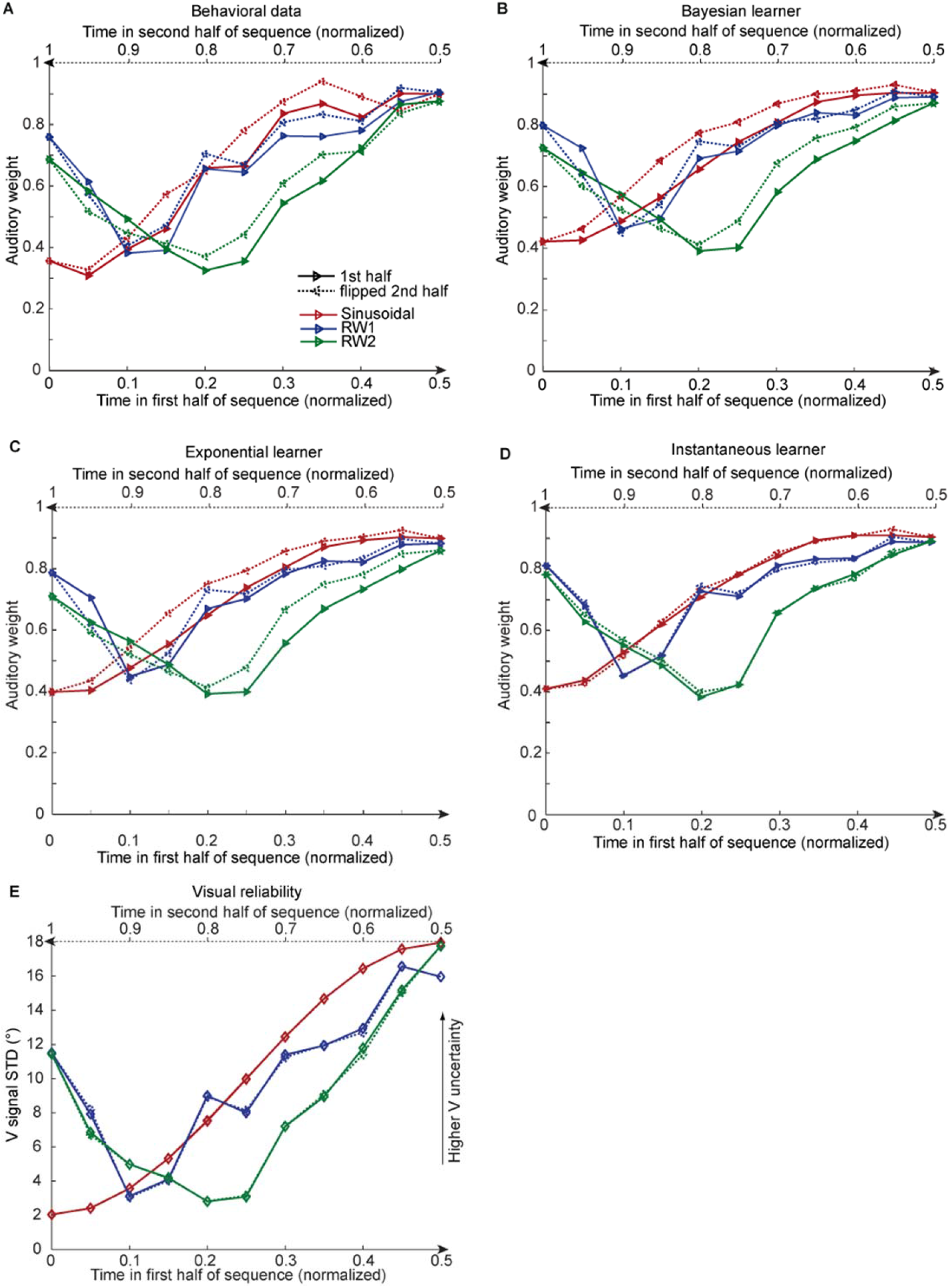
Observed and predicted relative auditory weights for continuous sequences of visual noise. Relative auditory weights w_A_ of the 1^st^ (solid) and the flipped 2^nd^ half (dashed) of a period (binned into 20 time bins) plotted as a function of the normalized time in the sinusoidal (red), the RW1 (blue) and the RW2 (green) sequences. Relative auditory weights were computed from auditory localization responses of human observers **(A)**, Bayesian **(B)**, exponential **(C)** or instantaneous **(D)** learning models. For comparison, the standard deviation of the visual signal is shown in **(E)**. Please note that all models were fitted to observers’ auditory localization responses (i.e. not the auditory weight w_A_).

**Figure 4.**
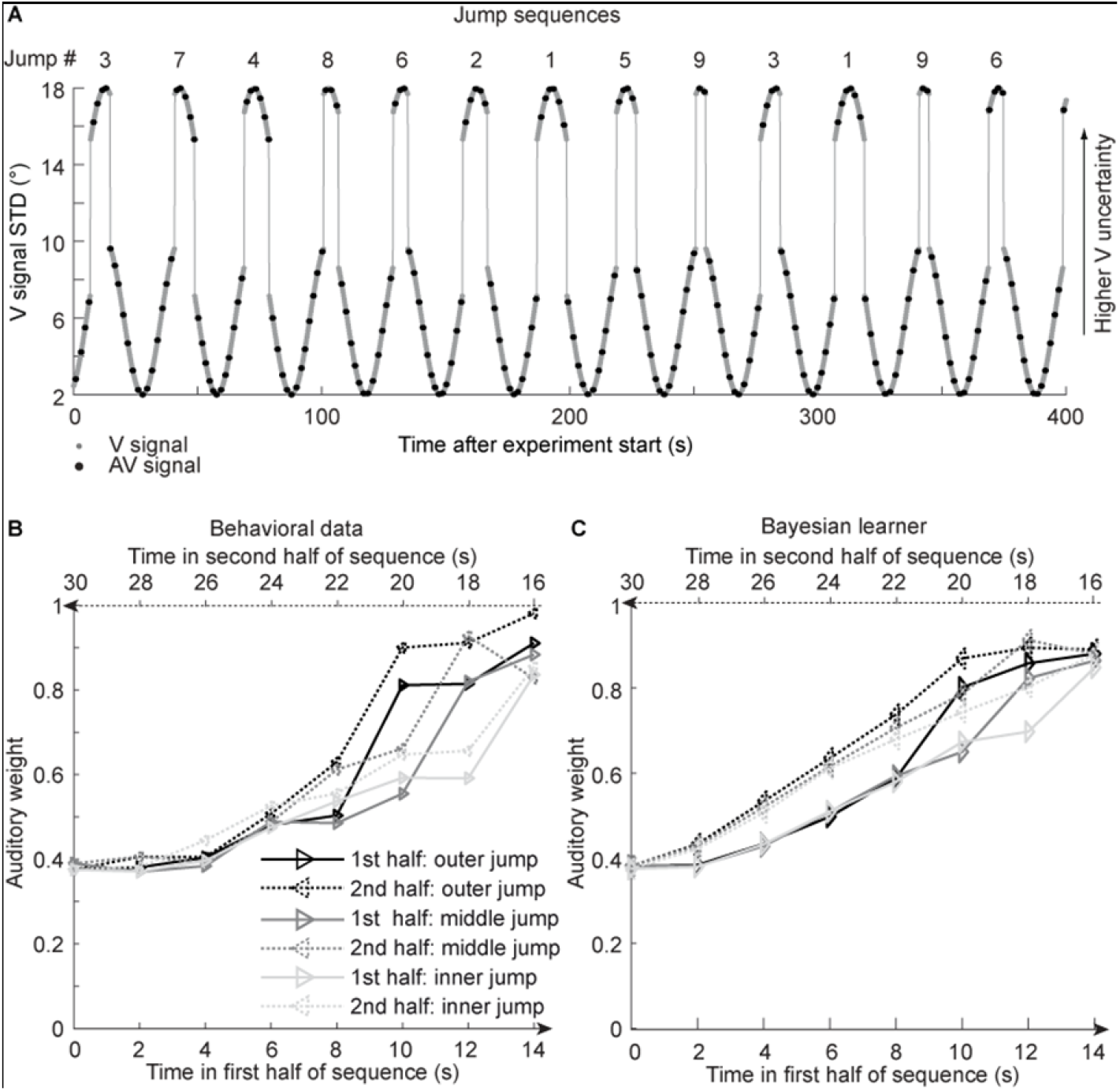
Time course of visual noise and relative auditory weights for sinusoidal sequence with intermittent jumps in visual noise. **(A)** The visual noise (i.e., STD of the cloud of dots, right ordinate) is displayed as a function of time. Each cycle included one abrupt increase and decrease in visual noise. The sequence of visual clouds was presented at 5 Hz while audiovisual (AV) signals (black dots) were interspersed with a SOA jittered between 1.4 and 2.8 s. **(B, C)** Relative auditory weights w_A_ of the 1^st^ (solid) and the flipped 2^nd^ half (dashed) of a period (binned into 15 time bins) plotted as a function of the time in the sinusoidal sequence with intermitted inner (light gray), middle (gray) and outer (dark gray) jumps. Relative auditory weights were computed from auditory localization responses of human observers **(B)** and the Bayesian learning model **(C).** Please note that all models were fitted to observers’ auditory localization responses (i.e. not the auditory weight w_A_).

We assigned physical visual noise (i.e., the cloud’s STD) and sound localization responses to 20 (resp. 15 for experiment 4) temporally adjacent bins in each period of the three sequences. Next, we quantified the auditory and visual influence on the perceived auditory location for each time bin based on the regression model: R_A_= L_A_* ß_A_ + L_V_* ß_V_ + const * ß + e with R_A_= Localization response; L_A_ or L_V_ = ‘true’ auditory or visual location; ß_A_ or ß_V_ = auditory or visual weight. The relative auditory weight was computed as w_A_ = ß_A_ / (ß_A_ + ß_V_). As expected, the auditory weight approximately tracked the visual cloud’s STD (Fig. 2). Yet, while the temporal evolution of the physical visual noise was designed to be symmetrical for each time period, we observed a temporal asymmetry for w_A_ in all three experiments. w_A_ was smaller for the 1^st^ half of each period, when visual noise increased, than the 2^nd^ half, when visual noise decreased over time (Fig. 3A). This impression was confirmed statistically in 2 (1^st^ vs. flipped 2^nd^ half) x 9 (time bins) repeated measures ANOVAs (Table 1) showing a significant main effect of 1^st^ versus flipped 2^nd^ period’s half for the sinusoidal (F(1, 24) = 12.162, p = 0.002, partial η^2^ = 0.336) and the RW1 sequence (F(1, 32) = 14.129, p < 0.001, partial η^2^ = 0.306). For the RW2 sequence, we observed a significant interaction (F(4.6, 82.9) = 3.385, p = 0.010, partial η^2^ = 0.158), because the visual noise did not change monotonously within each period half. Instead, monotonic increases and decreases in visual noise alternated at nearly the double frequency in RW2 as compared to RW1. The asymmetry in the auditory weights’ time course across the three experiments suggested that the visual noise in the past influenced observers’ current visual uncertainty estimate.

**Table 1.**
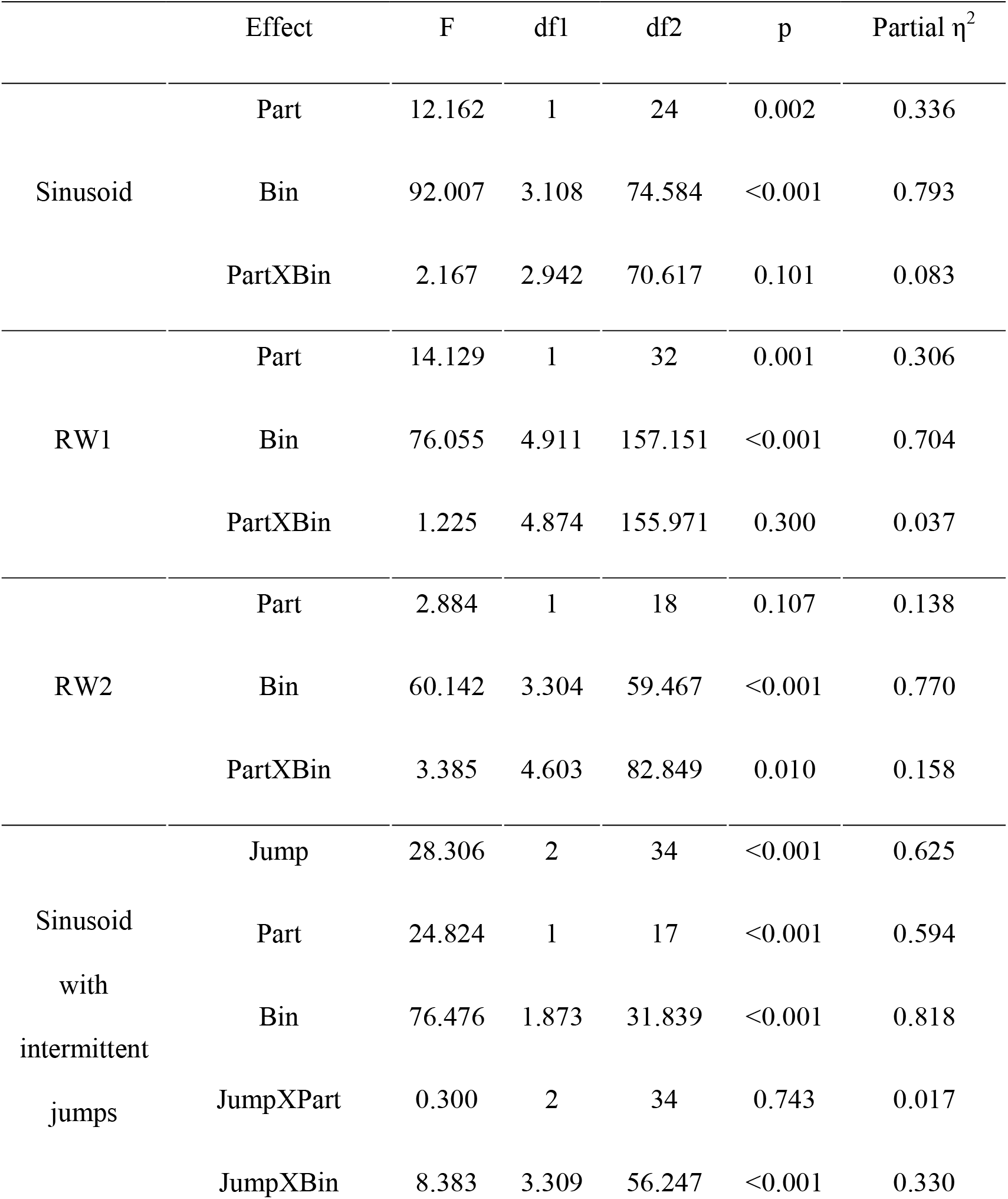
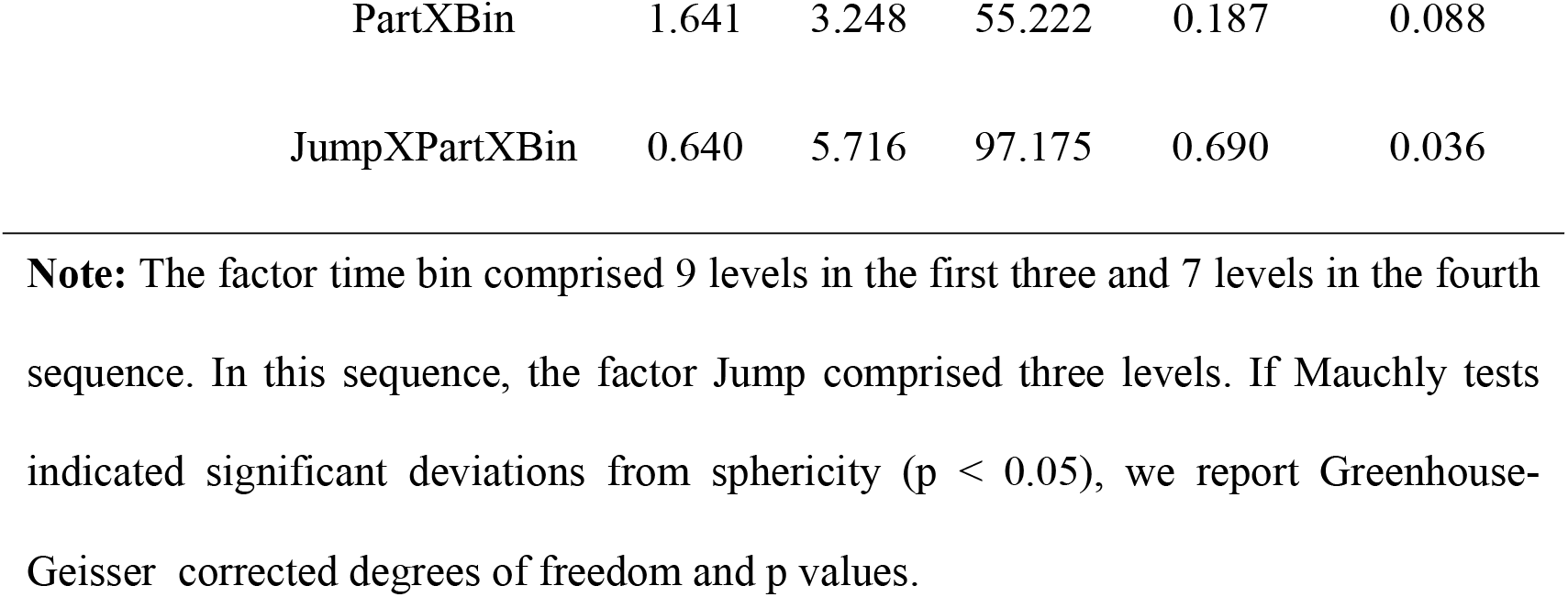
Analyses of the temporal asymmetry of the relative auditory weights across the four sequences of visual noise using repeated measures ANOVAs with the factors sequence part (1^st^ vs. flipped 2^nd^ half), time bin and jump position (only for the sinusoidal sequences with intermittant jumps).

We further tested this in a regression model where the relative auditory weights w_A_ for each of the 20 time bins were predicted by the visual STD in the current time bin and the difference in STD between the current and the previous bin. Indeed, both the current visual STD (p < 0.001 for all three sequences; Sinusoid: t(24)=15.767, Cohen’s d=3.153; RW1: t(32)= 15.907, Cohen’s d=2.769; RW2: t(18)=12.978, Cohen’s d=2.9773, two sided one-sample t test against zero) and the difference in STD between the current and the previous time bin (i.e. Sinusoid t(24)= −3.687, p = 0.001, Cohen’s d=−0.738; RW1 t(32)= −2.593, p = 0.014, Cohen’s d=−0.451; RW2 t(18)=−2.394, p = 0.028, Cohen’s d=−0.549) significantly predicted observers’ relative auditory weights. Collectively, these results suggest that observers’ visual uncertainty estimates (as indexed by the relative auditory weights w_A_) depend not only on the current sensory signal, but also on the recent history of the sensory noise.

To characterize how human observers use information from the past to estimate current sensory uncertainty, we compared three computational models that differed in how visual uncertainty is learnt over time: Model 1, the instantaneous learner, estimates visual uncertainty independently for each trial as assumed by current standard models. Model 2, the optimal Bayesian learner, estimates visual uncertainty by updating the prior uncertainty estimate obtained from past visual signals with the uncertainty estimate from the current signal. Model 3, the exponential learner, estimates visual uncertainty by exponentially discounting past uncertainty estimates. All three models account for observers’ uncertainty about whether auditory and visual signals were generated by common or independent sources by explicitly modeling the two potential causal structures^14^ underlying the audiovisual signals (n.b. only the model component pertaining to the ‘common cause’ case is shown in Fig. 1B, for the full model see Supplementary Fig. 1). Models were fit individually to observers’ data by sampling from the posterior over parameters for each observer (Table 2). We compared the three models using the Watanabe-Akaike information criterion (WAIC) as appropriate for evaluating model samples^15^ (i.e., a low WAIC indicates a better model, with a difference greater than 10 considered very strong evidence for a model): across observers, the Bayesian learner (WAIC_Bayes_ = 254158) outperformed the exponential learner (ΔWAIC = +61) and was substantially better than the instantaneous learner (ΔWAIC = +400). Further, two-sided non-parametric permutation tests at the group level showed that the WAIC-values for the Bayesian learner were significantly greater across observers in comparison to the instantaneous learner (p<0.001) with a small non-significant trend relative to the exponential learner (p~0.091). As shown in Figure 3 both, the Bayesian and the exponential learner, accurately reproduced the temporal asymmetry for the auditory weights across all three experiments.

**Table 2.** Model parameters (median) and relative WAIC values for the three candidate models in the four sequences of visual noise.

| Sequence | Model | $\sigma_A$ | $P_{\text{common}}$ | $\sigma_0$ | $\kappa$ or $\gamma$ | $\Delta\text{WAIC}$ |
| --- | --- | --- | --- | --- | --- | --- |
| Sinusoid | Instantaneous learner | 5.56 | 0.63 | 8.95 | - | 109.9 |
| | Bayesian learner | 5.64 | 0.65 | 9.03 | $\kappa$: 7.37 | 0 |
| | Exponential discounting | 5.62 | 0.64 | 9.02 | $\gamma$: 0.23 | 45.6 |
| RW1 | Instantaneous learner | 6.30 | 0.69 | 8.46 | - | 89.0 |
| | Bayesian learner | 6.29 | 0.72 | 8.68 | $\kappa$: 8.06 | 0 |
| | Exponential discounting | 6.26 | 0.70 | 8.75 | $\gamma$: 0.33 | 32.3 |
| RW2 | Instantaneous learner | 6.36 | 0.72 | 10.79 | - | 201.3 |
| | Bayesian learner | 6.49 | 0.78 | 10.9 | $\kappa$: 6.7 | 0 |
| | Exponential discounting | 6.46 | 0.73 | 11.0 | $\gamma$: 0.25 | 46.3 |
| Sinusoid with intermittent jumps | Instantaneous learner | 6.38 | 0.65 | 8.19 | - | 94.9 |
| | Bayesian learner | 6.45 | 0.68 | 8.26 | $\kappa$: 6.13 | 0 |
| | Exponential discounting | 6.43 | 0.67 | 8.20 | $\gamma$: 0.24 | 1.64 |

From the optimal Bayesian learner we inferred observers’ estimated rate of change in visual reliability (i.e. parameter 1/*κ*). The sinusoidal sequence was estimated to change at a faster pace (median κ = 7.4 across observers) than the RW1 sequence (median κ = 8.1), but slower than the RW2 sequence (median κ =6.7 for RW2 sequence) indicating that the Bayesian learner accurately inferred that visual reliability changed at different pace across the three continuous sequences (see legend of Fig. 2). Likewise, the learning rates 1-γ of the exponential learner accurately reflect the different rates of change across the sequences (*γ*: Sinusoid 0.23, RW1 0.33, RW2 0.25). Both the Bayesian and the exponential learner thus estimated a smaller rate of change for the RW1 than the sinusoidal sequence. Further, the learning rates of the exponential learner imply that observers gave the visual inputs presented 4.1 (Sinusoid), 5.4 (RW1) and 4.3 (RW2) seconds before the current stimulus 5% of the weight they assigned to the current visual input to estimate the visual reliability.

To further disambiguate between the Bayesian and the exponential learner, we designed a fourth experimental ‘jump sequence’ that introduced abrupt increases or decreases in in physical visual noise at three positions into the sinusoidal sequence (Fig. 4A). Using the same analysis approach as for experiments 1-3, we replicated the temporal asymmetry for the auditory weights (Fig 4B). For all three ‘jump positions’ w_A_ was significantly smaller for the 1^st^ half of each period, when visual noise increased, than the 2^nd^ half, when visual noise decreased over time. The 3 (jump positions) x 2 (1^st^ vs. flipped 2^nd^ half) x 7 (time bins) repeated measures ANOVA showed a significant main effect of 1^st^ versus flipped 2^nd^ period’s half (F(1,17) = 24.824, p < 0.001, partial η^2^ = 0.594), while this factor was not involved in any higher-order interactions (see Table 1). Further, in a regression model the current visual STD (t(17) = 11.655, p < 0.001, Cohen’s d = 2.747) and the difference between current and previous STD (t(17) = −4.768, p < 0.001, Cohen’s d = −1.124) significantly predicted the relative auditory weights. Thus, we replicated our finding that the visual noise in the past influenced observers’ current visual uncertainty estimate as indexed by the relative auditory weights w_A_.

Bayesian model comparison showed that both the Bayesian learner (WAIC_Bayes_ = 83796) and the exponential learner (WAIC_Bayes_ = 83798 substantially outperformed the instantaneous learner (ΔWAIC = +95, two-sided non-parametric permutation tests at the group level on the WAIC values: p < 0.001). However, consistent with our Bayesian model comparison results for the continuous sequences, the Bayesian learner did not provide a better explanation for observers’ responses than the exponential learner (ΔWAIC = +2, permutation test on the WAIC values: p=0.44, see Table 2, Fig. 4C and Supplementary Fig. 2A).

Across four experiments that used continuous and discontinuous sequences of visual noise, we have shown that the Bayesian learner outperforms the instantaneous learner. However, in none of the four experiments were we able to decide whether observers adapted to changes in visual noise according to a Bayesian or an exponential learner. The key feature that distinguishes between the Bayesian and the exponential learner is that only the Bayesian learner adapts dynamically based on its uncertainty about its visual reliability estimates. As a consequence, the Bayesian learner should adapt faster than the exponential learner to increases in physical visual noise (i.e. spread of the visual cloud) but slower to decreases in visual noise. From the Bayesian learner’s perspective, the faster learning for increases in visual noise emerges because it is unlikely that visual dots form a large spread cloud under the assumption that the true visual spread of the cloud is small. Conversely, the Bayesian learner will adapt more slowly to decreases in visual variance, because under the assumption of a visual cloud with a large spread visual dots may form a small cloud by chance.

Critically, the asymmetric differences in adaptation rate between the exponential and the Bayesian learner arise because the Bayesian learner adapts its visual uncertainty estimates influenced by its uncertainty about its visual reliability estimate. These differences will therefore be amplified if we increase observer’s uncertainty about its visual reliability estimate by reducing the number of dots of the visual cloud from 20 to 5 dots. Based on simulations, we therefore explored whether we could experimentally discriminate between the Bayesian and exponential learner using continuous sinusoidal or discontinuous ‘jump’ sequences with visual clouds of only 5 dots. For the two sequences, we simulated the sound localization responses of 12 observers based on the Bayesian learner model and fitted the Bayesian and exponential learner models to the responses of each simulated Bayesian observer. Figure 5 shows observers’ auditory weights indexing their estimated visual reliability across time that we obtained from the fitted responses of the Bayesian (blue) and the exponential learner (green). The simulations reveal the characteristic differences in how the Bayesian and the exponential learner adapt their visual uncertainty estimates to increases and decreases in visual noise. As expected, the Bayesian learner adapts its visual uncertainty estimates faster than the exponential learner to increases in visual noise, but slower to decreases in visual noise. Nevertheless, these differences are relatively small, so that the difference in mean log likelihood between the Bayesian and exponential learner is only −1.82 for the sinusoidal sequence and −2.74 for the jump sequence.

**Figure 5.**
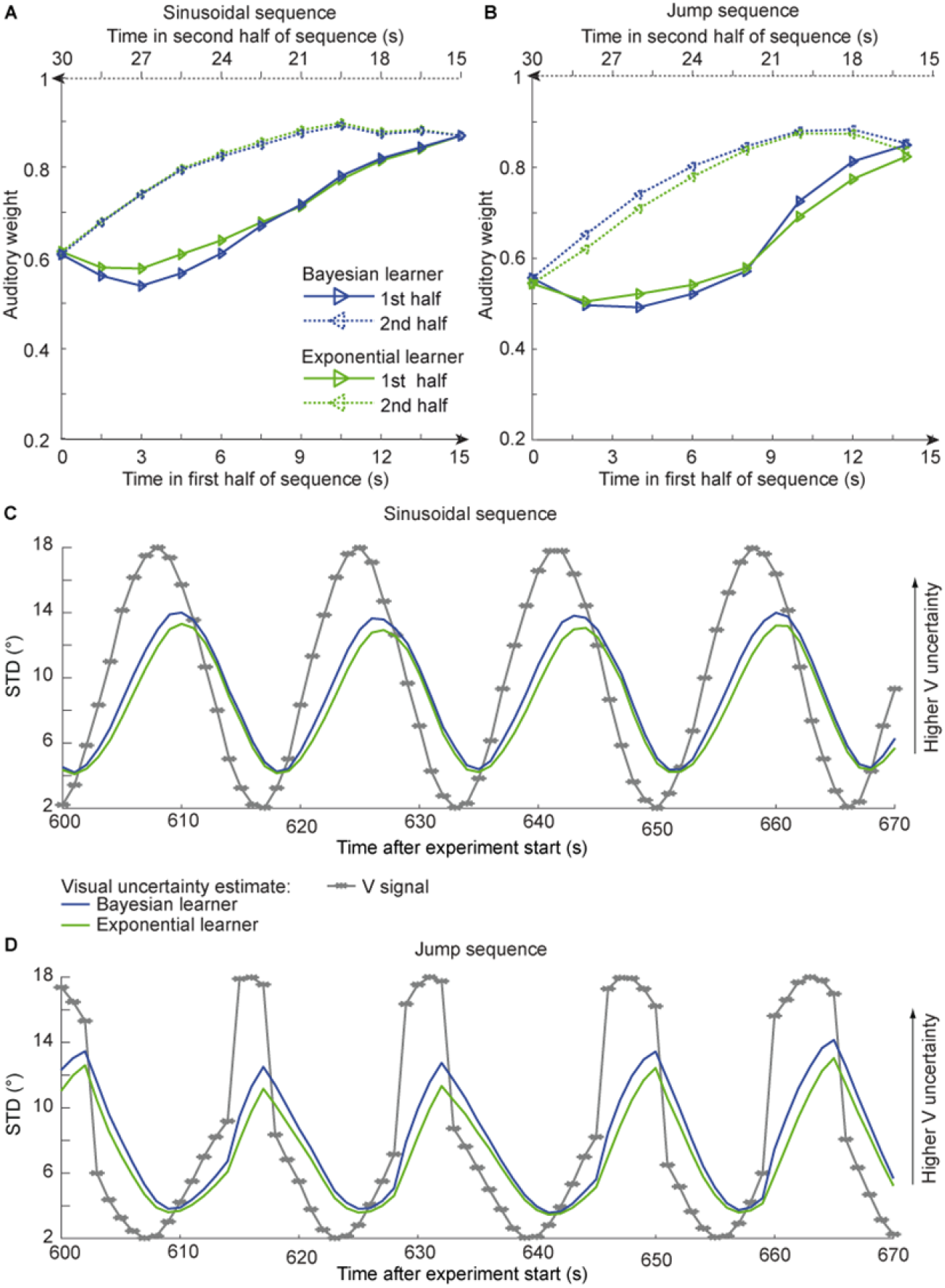
Time course of the relative auditory weights, the standard deviation of the visual cloud and the standard deviation of the visual uncertainty estimates. **(A)** Relative auditory weights w_A_ of the 1^st^ (solid) and the flipped 2^nd^ half (dashed) of a period (binned into 15 time bins) plotted as a function of the time in the sinusoidal sequence. Relative auditory weights were computed from the predicted auditory localization responses of the Bayesian (blue) or exponential (green) learning models fitted to the simulated localization responses of a Bayesian learner based on visual clouds of 5 dots. **(B)** Relative auditory weights w_A_ computed as in (A) for the sinusoidal sequence with intermitted jumps. Only the outer-most jump (dark grey in Fig. 4B/C and Supplementary Fig. 2) is shown. **(C, D)** Standard deviation (STD) of the visual cloud of 5 dots (grey) and the STD of observers’ visual uncertainty as estimated by the Bayesian (blue) and exponential (green) learners (that were fitted to the simulated localization responses of a Bayesian learner) as a function of time for the sinusoidal sequence (C) and in the sinusoidal sequence with intermitted jumps (D). Note that only an exemplary time course from 600-670 s after the experiment start is shown.

Next, we investigated whether our experiments successfully mimicked situations where observers benefit from integrating past and current information to estimate their sensory uncertainty. We compared the accuracy of the instantaneous, exponential and Bayesian learner’s visual uncertainty estimates in terms of their mean absolute deviation (in percentage) from the true variance. For Gaussian clouds of 20 dots, the instantaneous learner’s error in the visual uncertainty estimates of 21.7 % is reduced to 13.7 % and 14.9% for the exponential and Bayesian learners, respectively (with best fitted γ = 0.6, in the sinusoidal sequence). For Gaussian clouds composed of only 5 dots, the exponential and Bayesian learners even cut down the error by half (i.e. 46.8 % instantaneous learner, 29.5 % exponential learner, 23.9 % Bayesian learner, with best fitted γ = 0.7).

Collectively, these simulation results suggest that even in situations where observers benefit from combining past with current sensory inputs to obtain more precise uncertainty estimates, the exponential learner is a good approximation of the Bayesian learner, making it challenging to dissociate the two experimentally based on noisy human behavioural responses.

## Discussion

The results from our four experiments challenge classical models of perceptual inference where a perceptual interpretation is obtained using a likelihood that depends solely on the current sensory inputs^6^. These models implicitly assume that sensory uncertainty (i.e., likelihood variance) is instantaneously and independently accessed from the sensory signals on each trial based on initial calibration of the nervous system^16^. Most prominently, in the field of cue combination it is generally assumed that sensory signals are weighted by their uncertainties that are estimated only from the current sensory signals^5–7^ (but see^17,18^). By contrast, our results demonstrate that human observers integrate inputs weighted by uncertainties that are estimated jointly from past and current sensory signals. Across the three continuous and the one discontinuous jump sequences, observers’ current visual reliability estimates were influenced by visual inputs that were presented 4-5 s in the past albeit their influence amounted to only 5% of the current visual signals.

Critically, observers adapted their visual uncertainty estimates flexibly according to the rate of change in the visual noise across the experiments. As predicted by both Bayesian and exponential learning models, observers’ visual reliability estimates relied more strongly on past sensory inputs, when the visual noise changed more slowly across time. While observers did not explicitly notice that each of the four experiments was composed of repetition of temporally symmetric sequence components, we cannot fully exclude that observers may have implicitly learnt this underlying temporal structure. However, implicit or explicit knowledge of this repetitive sequence structure should have given observers the ability to predict and preempt future changes in visual reliability and therefore attenuated the temporal lag of the visual reliability estimates. Put differently, our experimental choice of repeating the same sequence component over and over again in the experiment cannot explain the influence of past signals on observers’ current reliability estimate, but should have reduced or even abolished it.

Importantly, the key feature that distinguishes the Bayesian from the exponential learner is how the two learners adapt to increases versus decreases in visual noise. Only the Bayesian learner represents and accounts for its uncertainty about its visual reliability estimates. As compared to the exponential learner, it should therefore adapt faster to increases but slower to decreases in visual noise. Our simulation results show this profile qualitatively, when the learner’s uncertainty about its visual reliability estimate is increased by reducing the number of dots (see Figure 5). But even for visual clouds of five dots, the differences in learning curves between the Bayesian and exponential learner are very small making it difficult to abjudicate between them given noisy observations. Unsurprisingly, therefore, Bayesian model comparison showed consistently across all four experiments that observers’ localization responses can be equally well explained by an optimal Bayesian and an exponential learner.

Collectively, our experimental and simulation results suggest that under circumstances where observers substantially benefit from combining past and current sensory inputs for estimating sensory uncertainty, optimal Bayesian learning can be approximated well by more simple heuristic strategies of exponential discounting where sensory weights are updated with a fixed learning rate that does not depend on observers’ uncertainty about their visual reliability estimate^3,10^. Future research will need to assess whether observers adapt their visual uncertainty estimates similarly if visual noise is manipulated via other methods such as stimulus luminance, duration or blur.

From the perspective of neural coding, our findings suggest that current theories of probabilistic population coding^12,19^ may need to be extended to accommodate additional influences of past experiences on neural representations of sensory uncertainties. Alternatively, the brain may compute sensory uncertainty using strategies of temporal sampling^20^.

In conclusion, to our knowledge, this is the first demonstration that human observers do not access sensory uncertainty instantaneously from the current sensory signals alone, but learn sensory uncertainty over time by combining past experiences and current sensory inputs as predicted by an optimal Bayesian learner or approximate strategies of exponential discounting. This influence of past signals on current sensory uncertainty estimates is likely to affect learning not only at slower timescales across trials (i.e. as shown in this study), but also at faster timescales of evidence accumulation within a trial^21^. While our research unravels the impact of prior sensory inputs on uncertainty estimation in a cue combination context, we expect that they reveal fundamental principles of how the human brain computes and encodes sensory uncertainty.

## Methods

### Participants

76 healthy volunteers participated in the study after giving written informed consent (40 female, mean age 25.3 years, range 18-52 years). All participants were naïve to the purpose of the study. All participants had normal or corrected-to normal vision and reported normal hearing. The study was approved by the human research review committee of the University of Tübingen.

### Stimuli

The visual spatial stimulus was a Gaussian cloud of twenty bright grey dots (0.56° diameter, vertical standard deviation 1.5°, luminance 106 cd/m^2^) presented on a dark grey background (luminance 62 cd/m^2^, i.e. 71% contrast). The auditory spatial cue was a burst of white noise with a 5ms on/off ramp. To create a virtual auditory spatial cue, the noise was convolved with spatially specific head-related transfer functions (HRTFs). The HRTFs were pseudo-individualized by matching participants’ head width, heights, depth and circumference to the anthropometry of subjects in the CIPIC database^22^. HRTFs from the available locations in the database were interpolated to the desired locations of the auditory cue.

### Experimental design and procedure

In a spatial ventriloquist paradigm, participants were presented with audiovisual spatial signals. Participants indicated the location of the sound by pressing one of 5 spatially corresponding buttons. The visual signal was a cloud of 20 dots sampled from a Gaussian. The visual clouds were re-displayed with variable horizontal standard deviations (see below) at a rate of 5 Hz (Fig. 1A). The cloud’s location mean was temporally independently resampled from five possible locations (−10°, −5°, 0°, 5°, 10°) at a stimulus onset asynchrony jittered between 1.4 and 2.8 s. In synchrony with the change in the cloud’s location, the dots changed their colour and a concurrent sound was presented. The location of the sound was sampled from ± 5° visual angle with respect to the mean of the visual cloud. Observers’ visual uncertainty estimate was quantified in terms of the relative weight the auditory signal on the perceived sound location. The change in the dot’s colour and the emission of the sound occurred in synchrony to enhance audiovisual binding.

### Continuous sinusoidal and random walk sequences

Critically, to manipulate visual noise over time, the cloud’s standard deviation changed according to i. a sinusoidal sequence, ii. a random walk sequence 1 or iii. a random walk sequence 2 (Fig. 2). In all sequences the horizontal standard deviation of the visual cloud spanned a range from 2-18°:

i. *Experiment1 - Sinusoidal sequence (Sinusoid):* A sinusoidal sequence was generated with a period of 30s. During the ~65 min of the experiment, each participant completed ~ 130 cycles of the sinusoidal sequence.
ii. *Experiment2 - Random walk sequence 1 (RW1):* First, we generated a random walk sequence of 60 s duration using a Markov chain with 76 discrete states and transition probabilities of stay (1/3), change to lower (1/3) or upper (1/3) adjacent states. To ensure that the random walk sequence segment starts and ends with the same value, this initial 60 s sequence segment was concatenated with its temporally reversed segment resulting in a RW sequence segment of 120 s duration. Each participant was presented with this 120s RW1 sequence approximately 32 times during the experiment.
iii. *Experiment3 - Random walk sequence 2 (RW2):* Likewise, we created a second random-walk sequence of 15 s duration using a Markov chain with only 38 possible states and transition probabilities similar to above. The 15 s sequence was concatenated with its temporally reversed version resulting in a 30 s sequence. The smoothness of this sequence segment was increased by filtering it (without phase shift) with a moving average of 250 ms. Each participant was presented with this sequence segment ~130 times.

Generally, a session of a Sinusoid, RW1 or RW2 sequence included 1676 trials. Because of experimental problems, four sessions included only 1128, 1143 or 1295 trials. Before the experimental trials, participants practiced the auditory localization task in 25 unimodal auditory trials, 25 audiovisual congruent trials with a single dot as visual spatial cue and 75 trials with stimuli as in the main experiment.

### Experiment4 - Sinusoidal sequence with intermittent changes in visual noise (sinusoidal jump sequence)

To dissociate the Bayesian learner from approximate exponential discounting, we designed a sinusoidal sequence (period = 30 s) with intermittent increases / decreases in visual variance (Fig. 4). As shown in Figure 4A, we inserted increases by 8° in visual STD at three levels of visual STD: 7.2°, 8.6°, 9.6° STD. Conversely, we inserted decreases by 8° in visual STD at 15.3°, 16.7°, 17.7° STD. We factorially combined these 3 (increases) x 3 (decreases) such that each sinewave cycle included exactly one sudden increase and decrease in visual STD (i.e., 9 jump types). Otherwise, the experimental paradigm and stimuli were identical to the continuous sinusoidal sequence described above. During the ~80 min of this experiment, each participant completed ~ 154 cycles of the sinusoidal sequence including 16-18 cycles for each of the 9 jump types.

This sinusoidal jump sequence aimed to maximize differences in adaptation rate for the Bayesian and exponential learner by introducing rapid increases or decreases in visual noise. While the exponential learner will weight past and present uncertainties throughout the entire sequence based on the same exponential function, the Bayesian learner will take into account the uncertainty about the visual variance and therefore adapt its visual variance estimate more slowly than the exponential learner for jumps from high to low visual uncertainty (see Fig. 5).

### Subject numbers and inclusion criteria

30 of the 76 subjects participated in the sinusoidal and the RW1 sequence session. Eight additional subjects participated only in the RW1 sequence session. 18 additional subjects participated in the RW2 sequence session. One participant completed all three continuous sequences. 20 subjects participated in the sinusoidal sequence with intermittent changes in visual uncertainty. In total, we collected data from 30 participants for the sinusoidal, 38 participants for the RW1, 19 participants for the RW2 and 20 participants for the sinusoidal jump sequence. From these samples, we excluded participants if their perceived sound location did not depend on the current visual reliability (i.e., inclusion criterion p < 0.05 in the linear regression; please note that this inclusion criterion is orthogonal to the question of whether participants’ visual uncertainty estimate depends on visual signals prior to the current trial). Thus, we excluded five participants of the sinusoidal and RW1 sequence and two participants from the sinusoidal jump sequence. Finally, we analysed data from 25 participants for the sinusoidal, 33 participants for the RW1, 19 participants for the RW2 and 18 participants for the sinusoidal jump sequence.

### Experimental setup

Audiovisual stimuli were presented using Psychtoolbox 3.09^23,24^ (www.psychtoolbox.org) running under Matlab R2010b (MathWorks) on a Windows machine (Microsoft XP 2002 SP2). Auditory stimuli were presented at ~75 dB SPL using headphones (Sennheiser HD 555). As visual stimuli required a large field of view, they were presented on a 30” LCD display (Dell UltraSharp 3007WFP). Participants were seated at a desk in front of the screen in a darkened booth, resting their head on an adjustable chin rest. The viewing distance was 27.5 cm. This setup resulted in a visual field of approx. 100°. Participants responded via a standard QWERTY keyboard. Participants used the buttons {i, 9, 0, −, =} with their right hand for localization responses.

### Data analysis

#### Continuous sinusoidal and random walk sequences

For each period of the three continuous sinusoidal and random walk sequences, we sorted the physical visual uncertainty (i.e., the cloud’s STD) and sound localization responses into 20 temporally adjacent bins. Next, we quantified the auditory and visual influence on the perceived auditory location for each time bin based on the regression model: R_A_= L_A_* ß_A_ + L_V_* ß_V_ + const * ß + e with R_A_= Localization response; L_A_ = ‘true’ auditory location; L_V_ = ‘true’ visual location; ß_A_ = auditory weight; ß_V_ = visual weight. The relative auditory weight was computed as w_A_ = ß_A_ / (ß_A_ + ß_V_) (Fig. 2A-C).

By design, the temporal evolution of the physical visual uncertainty (i.e., STD of the visual cloud) is symmetric for each period in the sinusoidal, RW1 and RW2 sequences. In other words, for physical visual noise the 1^st^ half and the flipped 2^nd^ half within a period are identical (Fig. 3E). Given this symmetry constraint, we evaluated the influence of past visual uncertainty on participants’ auditory weight w_A_ in terms of the difference between 1^st^ half and flipped 2^nd^ half. If prior visual noise affects current visual uncertainty estimates, we would expect smaller w_A_ for the 1^st^ half, when visual noise increased, than the 2^nd^ half, when visual noise decreased over time. We evaluated this statistically in 2 (1^st^ vs. flipped 2^nd^ half) x 9 (time bins, i.e. removing the bins at maximal and minimal visual noise values) repeated measures ANOVAs performed independently for the sinusoidal, RW1 and RW2 experiments (Table 1).

To further test whether the noise of past visual signals influenced observers’ current visual uncertainty estimate, we employed a regression model where the relative auditory weights were predicted by the visual STD in the current time bin and the difference in STD between the current and the previous time bin: w_A,t_= σ_V,t_ * ß_t_ + (σ_V,t_ – σ_V,t−1_)*ß_dt_ +const * ß+ e with w_A,t_= relative auditory weight in time bin i; σ_V,t_= visual STD in time bin i. To allow for generalization to the population level, the parameter estimates (ß_t,_ ß_dt_) for each subject were entered into two-sided one-sample t-tests at the between-subject random-effects level.

#### Sinusoidal sequence with intermittent changes in visual uncertainty

For each period of the sinusoidal sequence with intermittent changes, we sorted the physical visual uncertainty (i.e., the cloud’s STD) and sound localization responses into 15 temporally adjacent bins which were positioned to optimally capture the jumps in visual noise. For analysis of these sequences, we recombined the first and second halves of the 3 (increases at low, middle, high) x 3 (decreases at low, middle, high) sinewave cycles into three types of sinewave cycles such that both jumps were at low (= outer jump), middle (=middle jump) or high (= inner jump) visual uncertainty. This recombination makes the simplifying assumption that the jump position of the first half will have negligible effects on participants’ uncertainty estimates of the second half. As for the continuous sequences, we quantified the auditory and visual influence on the perceived auditory location for each time bin based on the regression model: R_A_= L_A_* ß_A_ + L_V_* ß_V_ + const * ß + e independently for the 15 temporally adjacent bins and computed the relative auditory weight w_A_. We statistically evaluated the influence of past visual noise on participants’ auditory weight on the w_A_ in terms of the difference between 1^st^ half and flipped 2^nd^ half using a 2 (1^st^ vs. flipped 2^nd^ half) x 7 (time bins) x 3 (jump: inner, middle, outer) repeated measures ANOVAs (Table 1).

### Computational Models (for continuous and discontinuous sequences)

To further characterize whether and how human observers use their uncertainty about previous visual signals to estimate their uncertainty of the current visual signal, we defined and compared three models where the visual reliability (λ_V_) was (1) estimated instantaneously for each trial (i.e., instantaneous learner), was updated via (2) Bayesian learning or (3) exponential discounting (i.e. exponential learner) (Supplementary Fig. 1).

In the following, we will first describe the generative model that accounts for the fact that visual uncertainty usually changes slowly across trials (i.e. time-dependent uncertainty changes) and auditory and visual signals can be generated by one common or two independent sources (i.e. causal structure). Starting from this generative model that combines causal inference and sensory uncertainty learning, we will describe the models for the instantaneous learner, the Bayesian learner and the exponential learner. Finally, we will explain how we account for participants’ internal noise and predict participants’ responses from each model (i.e. the experimenter’s uncertainty).

#### Generative model

On each trial *t,* the subject is presented with an auditory signal *A*_,*t*_, from a source *S*_*A,t*_, (see Supplementary Figure 1) together with a visual cloud of dots at time *t* arising from a source, *S*_,*Vt*_, drawn from a Normal distribution *S*_,*Vt*_ ~*N(0,* 1/λ_S_) with the spatial reliability (i.e., inverse of the spatial variance): λ_S_ = 1/*σ*_*s*_^2^. Critically, *S*_*A,t*_ and *S*_*V,t*_, can either be two independent sources (C = 2) or one common source (C=1): *S*_*A,t*_ = *S*_*V,t*_ = *S*_*t*_ ^14^.

We assume that the auditory signal is corrupted by noise, so that the internal signal is *A*_*t*_ ~ *N(S*_*A,t*_, 1/λ_A_). By contrast, the individual visual dots (presented at high visual contrast) are assumed to be uncorrupted by noise, but presented dispersed around the location *S*_*V,t*_ according to *V*_*i,t*_ ~ *N(U*_*t*_, 1/λ_V,t_), where *U*_*t*_ ~ *N(S*_*V,t*_, 1/λ_V,t_). The dispersion of the individual dots, 1/λ_V,t_, is assumed to be identical to the uncertainty about the visual mean, allowing subjects to use the dispersion as an estimate of the uncertainty about the visual mean.

The visual reliability of the visual cloud, λ_V,t_ = 1/σ_V,t_^2^, varies slowly at the re-display rate of 5 Hz according to a log random walk: log λ_V,t_ ~N(log λ_V,t−1_, 1/*κ*) with 1/*κ* being the variability of λ_V,t_ in log space. We also use this log random walk model to approximate learning in the four jump sequence (see ^25^).

The generative models of the instantaneous, Bayesian and exponential learners all account for the causal uncertainty by explicitly modeling the two potential causal structures. Yet, they differ in how they estimate the visual uncertainty on each trial, which we will describe in greater detail below.

#### Observer Inference

The instantaneous, Bayesian and exponential learners invert this (or slightly modified, see below) generative model during perceptual inference to compute the posterior probability of the auditory location, *S*_*A,t*_, given the observed *A*_*t*_ and *V*_*i,t*_. The observer selects a response based on the posterior using a subjective utility function which we assume to be the minimization of the squared error (*S*_*A,t*_ − *S*_*true*_)^2^. For all models, the estimate for the location of the auditory source is obtained by averaging the auditory estimates under the assumption of common and independent sources whether by their respective posterior probabilities (i.e. model averaging, see Supplementary Figure 1):

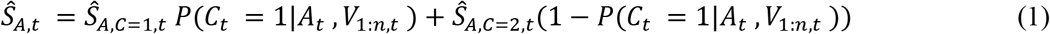

where 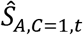 and 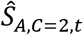 depend on the model (see below), and *P*(*C* = 1|*A*_*t*_, *V*_1:*n,t*_) is the posterior probability that the audio and visual stimuli originated from the same source according to Bayesian causal inference^14^.

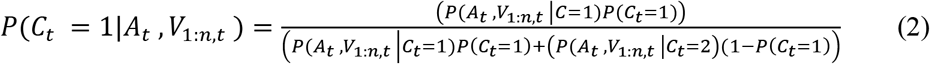

Finally, for all models we assume that the observer pushes the button associated with the position closest to 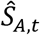. In the following, we describe the generative and inference models for the instantaneous, Bayesian and exponential learners. For the Bayesian learner, we focus selectively on the model component that assumes a common cause, C = 1 (for full derivation including both model components see Supplementary Material).

#### Model 1: Instantaneous learner

The instantaneous learning model ignores that the visual reliability (i.e., the inverse of visual uncertainty) of the current trial depends on the reliability of the previous trial. Instead, it estimates the visual reliability independently for each trial from the spread of the cloud of visual dots:

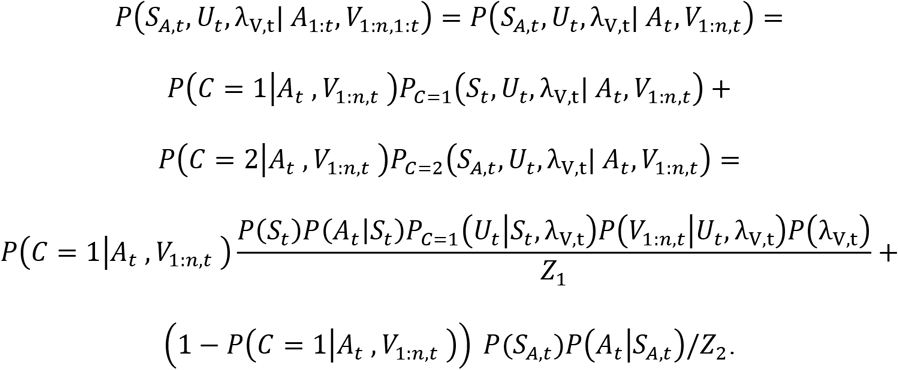

with *Z*_1_, *Z*_2_ as normalization constants.

Apart from *P*(*C* = 1|*A*_*t*_, *V*_*t*_), these terms are all normal distributions, while we assume in this model that *P*(λ_V,t_) is uninformative. Hence, visual reliability is computed from the sample variance: 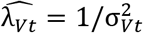 where 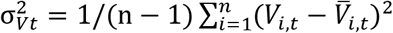 is the sample variance (and 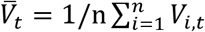 is the sample mean). The causal component estimates are given by:

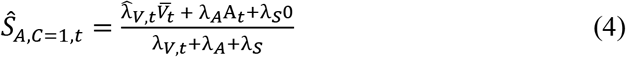

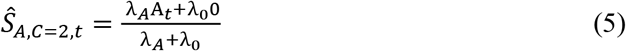

These two components are then combined based on the posterior probabilities of common and independent cause models (see equation 1). This model is functionally equivalent to a Bayesian causal inference model as described in Koerding et al. (2007)^14^, but with visual reliability computed directly from the sample variance rather than a fixed unknown parameter (which the experimenter estimates during model fitting).

#### Model 2: Bayesian learner

The Bayesian learner capitalizes on the slow changes in visual reliability across trials and combines past and current inputs to provide a more reliable estimate of visual reliability and hence auditory location. It computes the posterior probability based on all auditory and visual signals presented until time t (here only shown for C=1, see Supplementary Material for full derivation).

According to Bayes rule, the joint probability of all variables until time *t* can be written based on the generative model as:

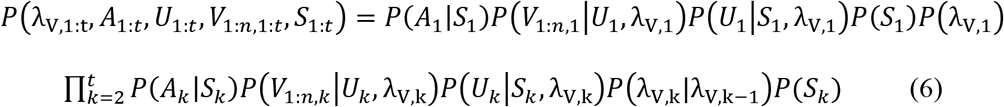

As above, the visual likelihood is given by the product of individual Normal distributions for each dot *i*: 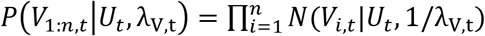, and *P*(*U*_*t*_|*S*_*t*_, λ_V,t_) = *N*(*U*_*t*_|*S*_*t*_, 1/λ_V,t_). The prior *P*(*S*_*t*_) is a Normal distribution *N*(*S*_*t*_|0, 1/λ_S_) and the auditory likelihood *P*(*A*_*t*,_|*S*_*t*_) is a Normal distribution *N*(*A*_*t*_|*S*_*t*_, 1/*λ*_*A*_). As described in the generative model, *P*(λ_V,k_|λ_V,k−1_) is given by log λ_V,t_ ~*N*(log λ_V,t−1_, 1/*κ*).

Importantly, only the visual reliability, λ_V,t_, is directly dependent on the previous trial (*P*(λ_V,k_, λ_V,k−1_) = *P*(λ_V,k_|λ_V,k−1_)*P*(λ_V,k−1_) ≠ *P*(λ_V,k_)*P*(λ_V,k−1_)). Because of the Markov property (i.e. λ_V,t_ depends only on λ_V,t−1_), the joint distribution for time *t* can be written as

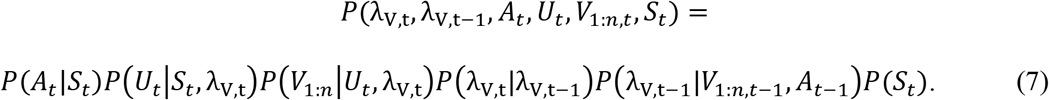

Hence, the joint posterior probability over location and visual reliability given a stream of auditory and visual inputs can be rewritten as:

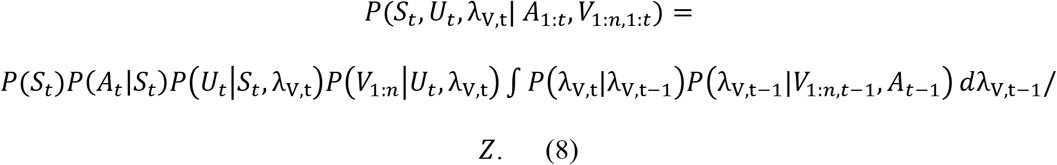

As this equation cannot be solved analytically, we obtain an approximate solution by factorizing the posterior in terms of the unknown variables (*S*_*t*_, *U*_*t*_, λ_V,t_) according to the method of variational Bayes^26^. For a full derivation of this approach, see Supplementary Material.

In short, a derivation is performed for the case of a common cause (C=1), as well as for the separate causes (C=2), and the two are then combined based on their relative posterior probability *P*(*C* = 1|*A*_*t*_, *V*_1:*n,t*_), as described above in equation 1.

#### Model 3: Exponential learner

Finally, the observer may approximate the full Bayesian inference of the Bayesian learner by a more simple heuristic strategy of exponential discounting. In the exponential discounting model, the observer learns the visual reliability by exponentially discounting past visual reliability estimates:

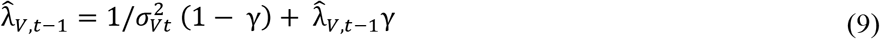

where 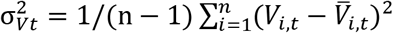 is the sample variance and 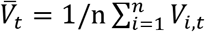 is the sample mean.

Similar to the optimal Bayesian learner (above), this observer model uses the past to compute the current reliability, but it does so based on a fixed learning rate 1 - γ. Computation is otherwise performed in accordance with models 1 and 2, equations 1–2 and 4–5.

#### Inference by the experimenter

From the observer’s viewpoint, this completes the inference process. However, from the experimenter’s viewpoint, the internal variable for the auditory stimulus, *A*_*t*_, is unknown and not directly under the experimenter’s control. To integrate out this unknown variable, we generated 1,000 samples of the internal auditory value for each trial from the generative process *A*_*t*_ ~ *N(S*_*A,t,true*_, *σ*_*A*_^*2*^), where *S*_*A,t,true*_ was the true location the auditory stimulus came from. For each value of A_t_, we obtained a single estimate 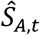 (as described above). To link these estimates with observers’ button response data, we assumed that subjects push the button associated with the position closest to 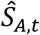. In this way, we obtained a histogram of responses for each subject and trial which provide the likelihood of the model parameters given a subject’s responses: *P*(*resp*_*t*_|*κ*, *σ*_*A*_, *P*_*common*_, *S*_*A,t,true*_, *S*_*V,t,true*_).

### Model estimation and comparison

Parameters for each model (for all models: σ_A_, P_common_= P(C=1), σ_0_, Bayesian learner: *κ*, exponential learner: γ) were fit for each individual subject by sampling using a symmetric proposal Metropolis-Hasting (MH) algorithm (with A_t_integrated out via sampling, see above). The MH algorithm iteratively draws samples *set*_*n*_ from a probability distribution through a variant of rejection sampling: if the likelihood of the parameter set is larger than the previous set, the new set is accepted, otherwise it is accepted with probability *L(model*|*set*_*n*_)/*L(model*|*set*_*n−1*_), where *L(resp*|*set*_*n*_)= ∏_*t*_ *P*(*resp*_*t*_|*κ*, *σ*_*A*_, *P*_*common*_, *S*_*A,t,true*_, *S*_*V,t,true*_) (for Bayesian learner). We sampled 4000 steps from 4 sampling chains with thinning (only using every 4^th^ sampling to avoid correlations in samples), giving a total of 4000 samples per subject data sets. Convergence was assessed through scale reduction (using criterion R<1.1^27^). Using sampling does not just provide a single parameter estimate for a data set (as when fitting maximum likelihood), but can instead be used to assess the uncertainty in estimation for the data set. The model code was implemented in Matlab (Mathworks, MA) and ran on two dual Xeon workstations. Each sample step, per subject data set, took 30 seconds on a single core (~42 hours per sampling chain).

Quantitative Bayesian model comparison of the three candidate models was based on the Watanabe-Akaike Information Criterion (WAIC) as an approximation to the out of sample cross validation^27^. For the comparison between models, we performed a non-parametric randomization test, comparing the observed difference in WAIC values with a null distribution where the labels of the models were randomly allocated (i.e. the sign of the WAIC difference between models for each subject were randomly permuted).

To qualitatively compare the localization responses given by the participants and the responses predicted by the instantaneous, Bayesian and exponential learner, we computed the auditory weight w_A_ from the predicted responses of the three models exactly as in the analysis for the behavioral data. For illustration, we show and compare the model’s w_A_ from the 1^st^ and the flipped 2^nd^ half of the periods for each of the four experiments (cf. Fig. 3, Fig. 4B/C and Supplementary Fig. 2).

### Parameter recovery

To test the validity of the models, we performed parameter recovery and were able to recover the generating values with a bias of all parameters smaller than 10 percent (for full details of bias and variance across parameters, see Supplementary Methods and Supplementary Tab. 1).

### Simulated localization responses

To further compare the Bayesian and exponential learner and assess whether they can be discriminated experimentally, we simulated the choices of 12 subjects for the continuous sinusoidal and sinusoidal jump sequence using the Bayesian learner model (parameters: σ_A_ =6 deg, κ=15, P_common_ =0.7 and σ_0_ =12 degrees). To increase observers’ uncertainty about their visual reliability estimates, we reduced the number of dots in the visual clouds from 20 to 5 dots where we ensured that the mean and variance of the 5 dots corresponded to the experimentally defined visual mean and variance. We then fitted the Bayesian learner and exponential learner models to each simulated data set (using the BADS toolbox for likelihood maximization^28^). The fitted parameters for the Bayesian model, *set*_*Bayes*_ were very close to the parameters used to generate observers’ simulated responses (sinusoidal sequence, fitted parameters: σ_A_ = 6.11°, κ = 17.5, P_common_ = 0.72 and σ_0_ = 12.4°; sinusoidal jump sequence, fitted parameters: σ_A_ = 6.08°, κ = 17.3, P_common_ = 0.71 and σ_0_ = 12.2°) – thereby providing a simple version of parameter recovery. The parameters of the exponential model, *set*_*Exp*_ (fitted to observers’ responses generated from the Bayesian model) were very similar to those of the Bayesian learner (sinusoidal sequence: σ_A_ = 5.99°, γ =0.70, P_common_ =0.61 and σ_0_ =12.0°, sinusoidal jump sequence: σ_A_ =6.06°, γ= 0.70, P_common_ =0.65 and σ_0_ =12.0°). Moreover, the fits to the simulated observers’ responses were very close for the two models (Figure 5), with mean log likelihood difference (*log(L(resp*|*set*_*Bayes*_)) – log(*L(resp*|*set*_*Exp*_))) =1.82 for the sinusoidal and 2.74 for the sinusoidal jump sequence (implying a slightly better fit for the Bayesian learner). Figure 5C and D show the timecourses of observers’ visual uncertainty (STD) as estimated by the Bayesian and exponential learners.

## Acknowledgements

This study was funded by the ERC (ERC-multsens, 309349) and the Max Planck Society and partially funded by the Deutsche Forschungsgemeinschaft (DFG; grant number RO 5587/1-1).

## Author contributions

TR, OS and UN conceived the experiment. TR and AF collected the data. UB, TR and UN analyzed the data. UB developed the computational models. UB, TR and UN wrote the manuscript.

## Competing interests

The authors report no conflict of interest.

## Notes

#### Summary of Updates

Updated version, with more analysis

