## Supplementary material for "Using the past to estimate sensory uncertainty"

#### Supplementary Methods

To test the validity of the Bayesian, exponential and instantaneous models, we performed parameter recovery by assessing the bias and variability of the parameters fitted to simulated data sets with respect to the true parameters used to generate the data.

For each model, we selected four different parameter sets (within a realistic range of values for parameters $\sigma_A$ = [6:12]°, $P_{common}$ = [0.7:0.9], $\sigma_0$ = [6:20]°, $\kappa$ = [5:20], $\gamma$ = [0.3:0.7]) and generated data sets of simulated observers for the RW2 sequence. We repeated this process 6 times (with different initial random seeds), creating a total of 24 simulated data sets for each model. We then fitted the Bayesian, exponential and instantaneous learner models to each simulated data set (using exactly the same fitting procedures as for observers' data in the experiments) resulting in 24 sets of best fitting parameters for each model.

In order to assess how well the fitting procedure recovers the generating parameters, we compared the fitted parameters to the 'true' parameters used to generate the data. Specifically, we assessed the parameter recovery in terms of bias and variability of the fitted parameters as follows: The recovered parameters' bias was computed as the signed deviation from the true generating value in percentage. As an example, if a data set was generated with a model parameter of 5, but the fitted (i.e. recovered) parameter was 4, we would compute a -20 % deviation. As a measure of the variability for the recovered parameters, we calculated the absolute (i.e. unsigned) deviation from the true generating values in percentage. As an example, a fitted value of 4, relative to a generating value of 5 would be a 20 percent absolute deviation. We report the median (and first and third quartile) across simulated data sets as a robust measure for this bias and variability..

Supplementary Table 1 shows the bias and variability of the recovered parameters for each model and each type of parameter. We report the median value as well as the boundary for the lower (25 percent) and upper (75 percent) quartile for both bias and variability. None of the parameters has a bias larger than 10 %, but the Bayesian learning parameter κ has a moderate variability of 32.1 %.

#### Supplementary figures

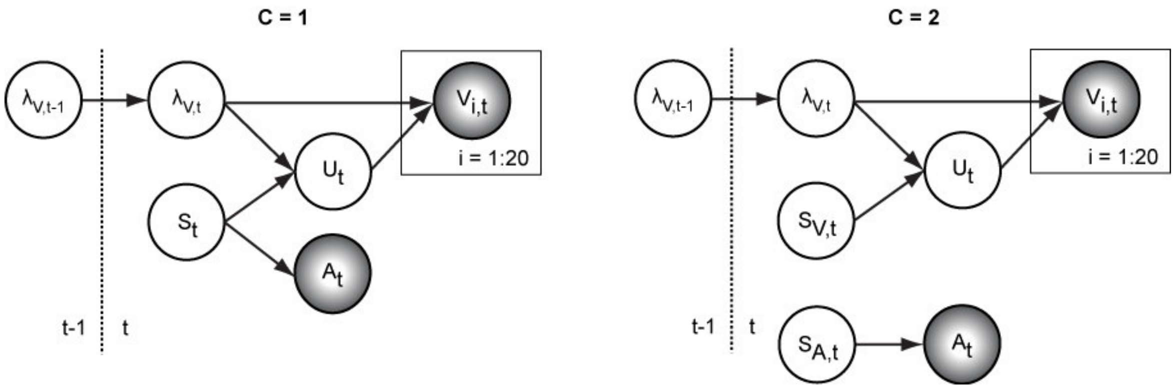

**Supplementary Figure 1. Generative model for the Bayesian learner.** The Bayesian Causal Inference model explicitly models whether auditory and visual signals are generated by one common (C=1) or two independent sources (C=2) (for further details see Koerding et al., 2007). We extend this Bayesian Causal Inference model into a Bayesian learning model by making the visual reliability ($\lambda_{V,t}$, i.e. the inverse of uncertainty or variance) of the current trial dependent on the previous trial.

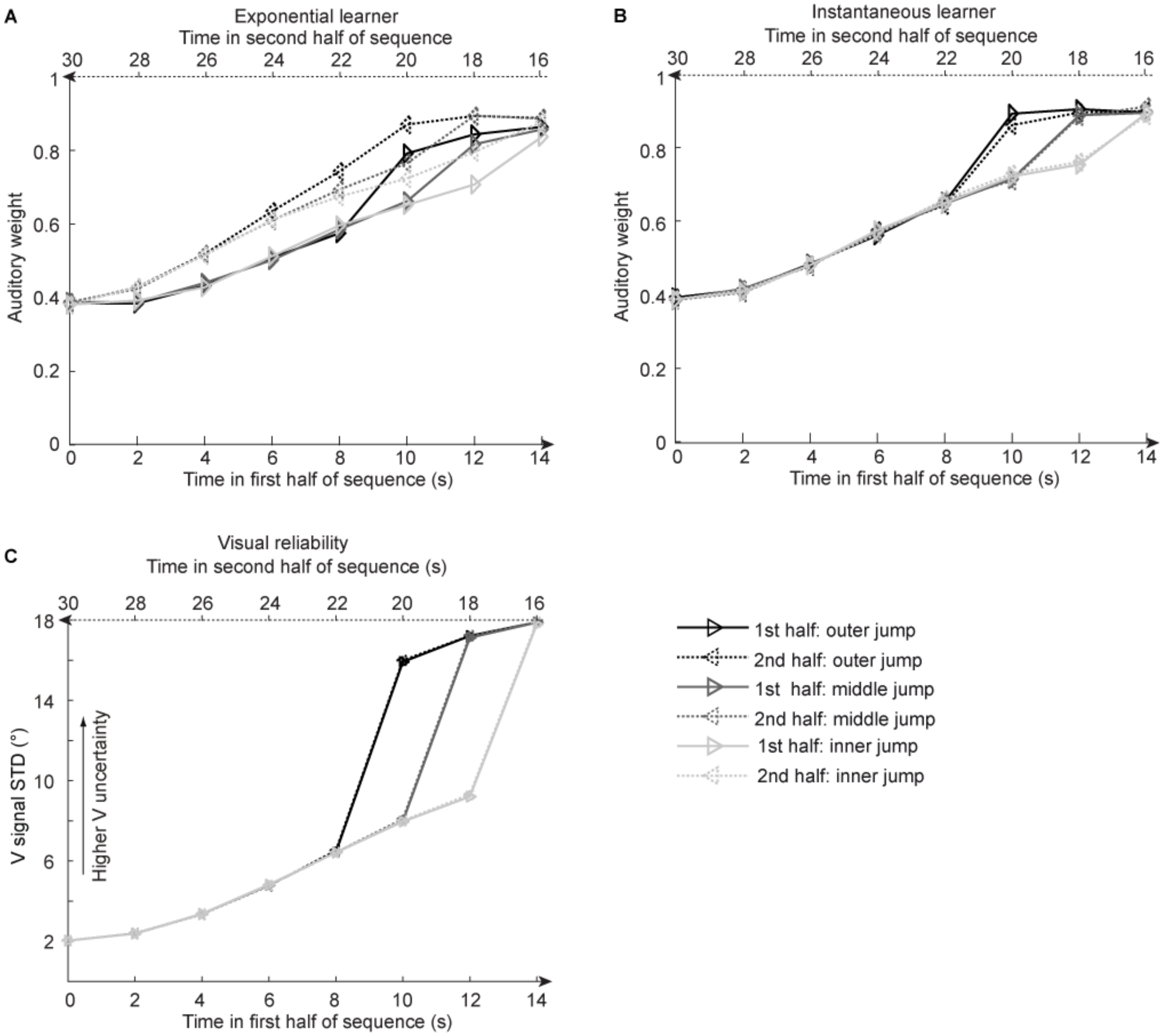

**Supplementary Figure 2. Time course of visual noise and relative auditory weights for the sinusoidal sequence with intermittent jumps in visual noise.** Relative auditory weights $w_A$ of the $1^{st}$ (solid) and the flipped $2^{nd}$ half (dashed) of a period (binned into 15 time bins) plotted as a function of the time in the sinusoidal sequence with intermitted inner (light gray), middle (gray) and outer (dark gray) jumps. Relative auditory weights were computed from auditory localization responses of exponential **(A)** or instantaneous **(B)** learning models. For comparison, the standard deviation of the visual signal is shown in **(C)**. Please note that all models were fitted to observers' auditory localization responses (i.e. not the auditory weight $w_A$).

#### Supplementary Table

**Supplementary Table 1. Percentage of bias and variability of model parameters from model recovery.**

| Model | Parameter | Bias | | | Variability | | |
|---|---|---|---|---|---|---|---|
| | | Q1 | Median | Q3 | Q1 | Median | Q3 |
| Instantaneous learner | $\sigma_A$ | -5.36 | -0.53 | 1.62 | 2.00 | 2.13 | 2.95 |
| | $P_{common}$ | -4.23 | -1.49 | 0.15 | 2.12 | 2.34 | 3.80 |
| | $\sigma_0$ | -3.12 | 1.81 | 5.01 | 2.62 | 3.42 | 9.45 |
| Bayesian learner | $\sigma_A$ | -6.70 | -0.17 | 2.15 | 2.36 | 3.22 | 5.52 |
| | $\kappa$ | -9.51 | 9.71 | 68.16 | 14.71 | 32.09 | 85.62 |
| | $P_{common}$ | -3.86 | -1.13 | 7.00 | 1.62 | 2.07 | 3.84 |
| | $\sigma_0$ | -2.26 | 2.68 | 4.97 | 3.00 | 4.45 | 6.79 |
| Exponential learner | $\sigma_A$ | -6.67 | -1.57 | 1.76 | 2.96 | 3.09 | 3.12 |
| | $\gamma$ | -5.43 | 4.43 | 8.02 | 6.17 | 7.11 | 12.69 |
| | $P_{common}$ | -4.92 | -2.23 | 0.18 | 2.63 | 3.28 | 4.47 |
| | $\sigma_0$ | -3.5 | 2.13 | 5.86 | 2.68 | 4.24 | 6.95 |

Note: The bias is computed by the percentage deviation from the true generating value. Variability is computed by the percentage absolute deviation from the true generating value. Q1 = first quartile; Q3 = third quartile

### Using the past to estimate sensory uncertainty - Supplementary Online Material

Ulrik Beierholm*,Tim Rohe*, Ambra Ferrari, Oliver Stegle, Uta Noppeney

This document describes a Variational Bayes approximation to inference on a generative model that allows for two possible ways that stimuli data was generated (thus allowing subjects to perform causal inference).

Section 1) describes the full generative model for both a single and two sources, section 2) explains how an optimal observer can perform inference within either sub-model, through a variational Bayes approximation to the posteriors and section 3) shows how to calculate the model likelihood for either sub-model, as necessary for combining the two sub-models.

Section 4) finally describes how the results for each causal model are combined into a single posterior.

**4 Putting it all together** **23**

### 1 Generative model

The model presented here is an extension of the Causal Inference model of Kording et al. 2007, with the reliability of the visual signal assumed to be changing smoothly over trials according to a random walk. In the case where the visual reliability is constant the model approximates the original Causal Inference model.

In this model (figure 1), at each stimulus presentation, $t$, subjects assume that the visual dots at positions $V_{i,t}$ and auditory stimulus at $A_t$, are generated through either of two causal models ($C_t = 1$ or $C_t = 2$) with fixed prior probabilities:

$$C_t \sim Bernoulli(pcommon) \tag{1}$$

If $C_t = 1$, (single source, $S_t$, leading to forced fusion)

$$S_t \sim \mathcal{N}(S_t; \mu_0, \sigma_0^2) \tag{2}$$
$$A_t \sim \mathcal{N}(A_t; S_t, \sigma_A^2) \tag{3}$$
$$U_t \sim \mathcal{N}(U_t; S_t, 1/\lambda_{V,t}) \tag{4}$$
$$V_{i,t} \sim \mathcal{N}(V_{i,t}; U_t, 1/\lambda_{V,t}) \tag{5}$$
$$\log \lambda_{V,t} \sim \mathcal{N}(\log(\lambda_{V,t}); \log(\lambda_{V,t-1}), k) \tag{6}$$

If $C_t = 2$ (independent sources, $S_{A,t}$ and $S_{V,t}$ )

$$S_{A,t} \sim \mathcal{N}(S_{A,t}; \mu_0, \sigma_0^2) \tag{7}$$
$$A_t \sim \mathcal{N}(A_t; S_{A,t}, \sigma_A^2) \tag{8}$$
$$S_{V,t} \sim \mathcal{N}(S_{V,t}; \mu_0, \sigma_0^2) \tag{9}$$
$$U_t \sim \mathcal{N}(U_t; S_{V,t}, 1/\lambda_{V,t}) \tag{10}$$
$$V_{i,t} \sim \mathcal{N}(V_{i,t}; U_t, 1/\lambda_{V,t}) \tag{11}$$
$$\log \lambda_{V,t} \sim \mathcal{N}(\log(\lambda_{V,t}); \log(\lambda_{V,t-1}), k) \tag{12}$$

The intermediate variable $U_t$ means that the mean of the visual dots is not located at the true source ($S_t$ or $S_{V,t}$), but normally distributed around it. Note that for $C_t = 1$ we can explicitly write $S_{A,t} = S_{V,t} = S_t$.

For simplicity we assume that $\mu_0 = 0$, i.e. the prior mean is located at the horizontal center. The auditory standard deviation $\sigma_A$, the prior probability of a single cause, $P_{common}$ and the prior standard deviation, $\sigma_0$ , are fixed individually for each subject (see main text for fitting procedure).

In the following we will simplify the notation by referring to $P(*|C_t = 1)$ by $P_1(*)$ and $P(*|C_t = 2)$ by $P_2(*)$.

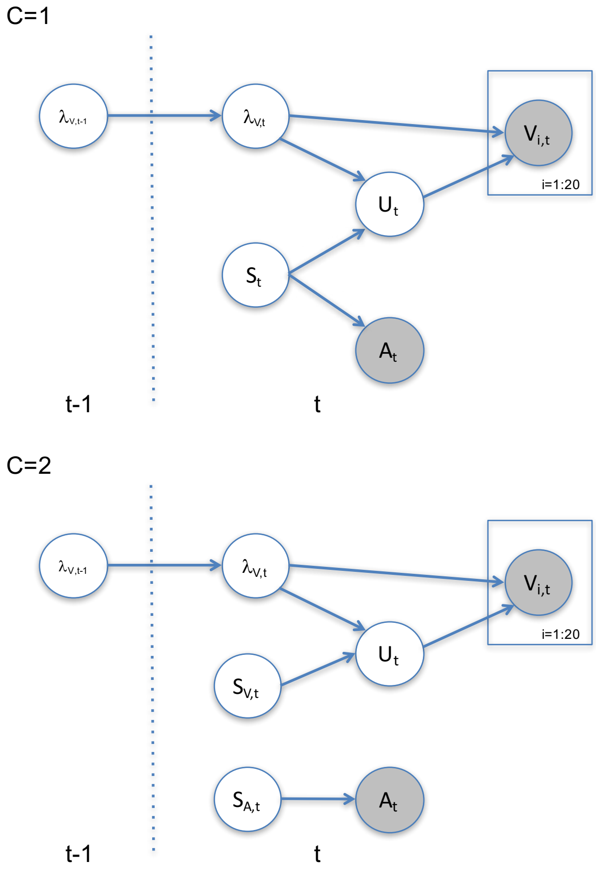

[h]

Figure 1: Generative model, for one (C=1) or two sources (C=2)

#### 2 Posterior

The full posterior over the latent variables in the model up until time $t$ is

$$P(S_{V,1:t}, S_{A,1:t},, \lambda_{1:t}, U_{1:t}, C_{1:t}|A_{1:t}, V_{1:N,1:t}) =$$
$$P(S_{V,1}, S_{A,1}, \lambda_{V,1}, U_1, C_1|A_1, V_{1:N,1}) \prod_{j=1}^{t} P(S_{V,j}, S_{V,j}, \lambda_{V,j}, U_j, C_j|A_{1:j}, V_{1:N,1:j}, \lambda_{V,j-1}) \tag{13}$$

Recursively we can write

$$P(S_{V,t}, S_{A,t}, \lambda_{V,t}, U_t, C_t|A_{1:t}, V_{1:N,1:t}) = \int P(S_{V,t}, S_{A,t}, \lambda_{V,t}, U_t, C_t|A_{1:t}, V_{1:N,1:t}, \lambda_{V,t-1})$$
$$P(S_{V,t-1}, S_{A,t-1}, U_{t-1}, C_t, \lambda_{V,1:t-1}|A_{1:t-1}, V_{1:N,1:t-1}) dS_{V,t-1} dS_{A,t} dU_{t-1} d\lambda_{V,1:t-1} \tag{14}$$

$$P(S_{V,t}, S_{A,t}, \lambda_{V,t}, U_t, C_t|A_{1:t}, V_{1:N,1:t}) \propto P(C_t)P(S_{A,t}, S_{V,t})P(A_t|S_{A,t})$$
$$P(V_{1:N,t}|U_t, \lambda_{V,t})P(U_t|S_{V,t},, \lambda_{V,t})$$
$$\int P(\lambda_{V,t}|\lambda_{V,t-1})P(\lambda_{V,1:t-1}|A_{1:t-1}, V_{1:N,1:t-1}) d\lambda_{V,1:t-1}/Z \tag{15}$$

where $P(S_{A,t}, S_{V,t})$ obviously depends on $C_t$ through the generative model (figure 1).

If we marginalise over the latent $C_t$:

$$P(S_{V,t}, S_{A,t}, \lambda_{V,t}, U_t|A_{1:t}, V_{1:N,1:t}) =$$
$$P(S_{V,t}, S_{A,t}, \lambda_{V,t}, U_t|A_{1:t}, V_{1:N,1:t}, C_t = 1)P(C_t = 1|A_{1:t}, V_{1:N,1:t})$$
$$+ P(S_{V,t}, S_{A,t}, \lambda_{V,t}, U_t|A_{1:t}, V_{1:N,1:t}, C_t = 2)P(C_t = 2|A_{1:t}, V_{1:N,1:t}) \tag{16}$$

At this point it should be clear that the posterior is a mixture of the forced fusion and independent solutions, with the mixture determined by the posterior probability of either model generating the data:

$$P(C_t = 1|A_{1:t}, V_{1:N,1:t}) = \frac{P(A_t, V_{1:N,t}|C_t = 1)P(C_t = 1)}{P(A_t, V_{1:N,t}|C_t = 1)P(C_t = 1) + P(A_t, V_{1:N,t}|C_t = 2)P(C_t = 2)} \tag{17}$$

To evaluate this we need to calculate the marginal model evidence, $P(A_t, V_{1:N,t}|C_t)$, for either model, see the later section.

##### 2.1 Posterior for C=1

The full posterior over the latent variables in the single source sub-model is

$$P_1(S_{1:t}, \lambda_{1:t}, U_{1:t}|A_{1:t}, V_{1:N,1:t}) = P(S_1, \lambda_{V,1}, U_1|A_1, V_{1:N,1:t}) \prod_{i=1}^{t} P(S_i, \lambda_{V,i}, U_i|A_i, V_i, \lambda_{V,i-1}) \tag{18}$$

Recursively we can write

$$P_1(S_t, \lambda_{V,t}, U_t|A_{1:t}, V_{1:N,1:t}) = \int P_1(S_t, \lambda_{V,t}, U_t|A_{1:t}, V_{1:N,1:t}, \lambda_{V,t-1}) \\ P_1(S_{t-1}, \lambda_{V,1:t-1}, U_{t-1}|A_{1:t-1}, V_{1:N,1:t-1}) dS_{t-1} dU_{t-1} d\lambda_{V,1:t-1} \quad (19)$$

$$P_1(S_t, \lambda_{V,t}, U_t|A_{1:t}, V_{1:N,1:t}) = \int P_1(S_t, \lambda_{V,t}, U_t|A_{1:t}, V_{1:N,1:t}, \lambda_{V,t-1}) \\ P(\lambda_{V,1:t-1}|A_{1:t}, V_{1:N,1:t-1}) d\lambda_{V,1:t-1} \quad (20)$$

$$P_1(S_t, \lambda_{V,t}, U_t|A_{1:t}, V_{1:N,1:t}) \propto P(S_t)P(A_t|S_t)P(U_t|S_t, \lambda_{V,t})P(V_{1:N,t}|U_t, \lambda_{V,t}) \\ \int P(\lambda_{V,t}|\lambda_{V,t-1})P(\lambda_{V,1:t-1}|A_{1:t-1}, V_{1:N,1:t-1}) d\lambda_{V,1:t-1}/Z \quad (21)$$

As we will see later it is convenient to use a change of parameters

$$\theta_t = \log(\lambda_{V,t}) \quad (22)$$

allowing us to rewrite

$$P_1(S_t, \theta_t, U_t|A_{1:t}, V_{1:N,1:t}) \propto P(S_t)P(A_t|S_t)P(U_t|S_t, \theta_t)P(V_{1:N,t}|U_t, \theta_t) \\ \int P(\theta_t|\theta_{V,t-1})P(\theta_{V,1:t-1}|A_{1:t-1}, V_{1:N,1:t-1}) d\theta_{V,1:t-1}/Z \quad (23)$$

where

$$P(U_t|S_t, \theta_t) = \mathcal{N}(U_t; S_t, 1/\exp(\theta_t)) \quad (24)$$

$$P(V_{1:N,t}|U_t, \theta_t) = \prod_n \mathcal{N}(U_t; V_{n,t}, 1/\exp(\theta_t)) \quad (25)$$

$$P(\theta_t|\theta_{V,t-1}) = \mathcal{N}(\theta_t; \theta_{V,t-1}, 1/\kappa) \quad (26)$$

We will assume that

$$P(\theta_{V,1:t-1}|A_{1:t-1}, V_{1:N,1:t-1}) \quad (27)$$

can be approximated by a Normal distribution (see below), thus allowing us to write

$$\int P(\theta_t|\theta_{V,t-1})P(\theta_{V,1:t-1}|A_{1:t-1}, V_{1:N,1:t-1}) d\theta_{V,1:t-1} = \mathcal{N}(\theta_t|\theta_{V,t-1}, 1/\kappa'_t) \quad (28)$$

where $1/\kappa'_t = 1/\kappa + 1/\tau_{\theta,t-1}$ (due to properties of convolution of two Normal distributions).

The log-posterior (to be used for a variational approximation) is now

$$\log P_1(S_t, \lambda_{V,t}, U_t|A_{1:t}, V_{1:N,1:t}) \propto -\lambda_S S_t^2/2 \\ - \lambda_A(A_t - S_t)^2/2 \\ - \lambda_{V,t}(U_t - S_t)^2/2 + \log \lambda_{V,t}/2 \\ - \lambda_{V,t}\sum_i^N (U_t - V_{i,t})^2/2 + (N/2)\log \lambda_{V,t} \\ - \kappa'(\theta_t - \theta_{V,t-1})^2/2 \quad (29)$$

#### 2.2 Variational Bayes approximation for C=1

We will now approximate the log-posterior with variational Bayes by factorization:

$$P_1(S_t, \theta_t, U_t|A_t, V_{1:N,t}) \approx q_1(S_t, U_t, \theta_t)) = q_1(S_t) * q_1(U_t) * q_1(\theta_t) \quad (30)$$

### 2.2.1 $q_1(S_t)$

For $q_1(S_t)$

$$\log q_1(S_t) \propto -\lambda_S S_t^2/2 - \lambda_A(A_t - S_t)^2/2 - E_\theta(\exp(\theta_t))E_U((U_t - S_t)^2)/2 \quad (31)$$

where $E_Y(X)$ signifies the expectation of X over the distribution of Y: $E_Y(X) = \int P(Y)XdY$.

Using $E_U((U_t - S_t)^2) = E_U(U_t^2 + S_t^2 - 2U_tS_t) = E_U(U_t^2) + S_t^2 - 2S_tE_U(U_t) = (S_t^2 - S_tE_U(U_t)*2 + E_U(U_t)^2) - E_U(U_t)^2 + E_U(U_t^2) = (S_t - E_U(U_t))^2 - E_U(U_t)^2 + E_U(U_t^2)$, where the last two terms do not depend on $S_t$ (and thus can be discarded) we can rewrite the last term:

$$\log q_1(S_t) \propto -\lambda_S S_t^2/2 - \lambda_A(A_t - S_t)^2/2 - E_\theta(\exp(\theta_t))(S_t - E_U(U_t))^2/2 \quad (32)$$

### 2.2.2 $q_1(U_t)$

For $q_1(U_t)$

$$\log q_1(U_t) \propto -E_\theta(\exp(\theta_t))/2\sum_i (U_t - V_{i,t})^2 - E_\theta(\exp(\theta_t))E_S((U_t - S_t)^2)/2 \quad (33)$$

Here we use the same trick

$$\log q_1(U_t) \propto -E_{\theta_t}(\exp(\theta_t))/2\sum_i (U_t - V_{i,t})^2 - E_\theta(\exp(\theta_t))(U_t - E_S(S_t))^2)/2 \quad (34)$$

###### 2.2.3 $q_1(\theta)$

For $q_1(\theta)$

$$\begin{aligned}\log q_1(\theta) = &-\exp\theta_t E_{U,S}((U_t - S_t)^2)/2 \\ &-\exp\theta_t E_U(\sum_i^N (U_t - V_{i,t})^2)/2 \\ &+(N+1)\log(\exp(\theta))/2 - \kappa'(\theta_t - \theta_{t-1})^2/2 \end{aligned} \quad (35)$$

###### 2.2.4 Simplifying $q_1(S_t)$ and $\log q_1(U_t)$

Inspecting $\log q_1(S_t)$ and $\log q_1(U_t)$ we can see that both $q_1(S_t)$ and $q_1(U_t)$ are products of Normal distributions, and thus themselves Normal distributed

$$q_1(S_t) \sim \mathcal{N}(S_t; \mu_{S,t}, 1/\tau_{S,t}) \quad (36)$$

and

$$q_1(U_t) \sim \mathcal{N}(U_t; \mu_{U,t}, 1/\tau_{U,t}) \quad (37)$$

where

$$\mu_{S,t} = (\lambda_S * 0 + \lambda_A A_t + E(\exp\theta_t)\mu_{U,t})/\tau_{S,t} \quad (38)$$

$$\tau_{S,t} = \lambda_S + \lambda_A + E(\exp(\theta_t)) \quad (39)$$

$$\mu_{U,t} = (E_\theta(\exp(\theta_t))\sum_i^N V_{i,t} + E_\theta(\exp(\theta_t))\mu_{S,t})/\tau_{U,t} \quad (40)$$

$$\tau_{U,t} = (N+1) * E_\theta(\exp(\theta_t)) \quad (41)$$

Note that $E_S(S_t) \approx \int q_1(S_t)S_t dS_t = \mu_{S,t}$ and $E_U(U_t) \approx \int q_1(U_t)U_t dU_t = \mu_{U,t}$

###### 2.2.5 Simplifying $q_1(\theta_t)$

Regarding $q_1(\theta_t)$ we can expand a little using that

$$\begin{aligned}E_{U,S}((U_t - S_t)^2) &= E_{U,S}((U_t^2 + S_t^2 - 2S_tU_t)) \\ &= E_{U,S}(U_t^2) + E(S_t^2) - 2E(S_t)E(U_t) \\ &= \mu_U^2 + 1/\tau_U + \mu_S^2 + 1/\tau_S - 2\mu_S\mu_U \\ &= (\mu_U - \mu_S)^2 + 1/\tau_U + 1/\tau_S \end{aligned} \quad (42)$$

(using that $E(X^2) = \mu^2 + 1/\tau$ for a normal distribution $\mathcal{N}(X; \mu, 1/\tau)$) and

$$E_U(\sum_i^N (U_t - V_{i,t})^2) = E_U(\sum_i^N (U_t^2 + V_{i,t}^2 - 2U_tV_{i,t}))$$
$$= E_U(\sum_i^N (U_t^2) + \sum_i^N (V_{i,t}^2) - 2\sum_i^N (U_tV_{i,t})$$
$$= E_U(N * (U_t^2) + \sum_i^N (V_{i,t}^2) - 2U_t \sum_i^N V_{i,t})$$
$$= N * (\mu_U^2 + 1/\tau_U) + \sum_i^N (V_{i,t}^2) - 2\mu_U \sum_i^N V_{i,t}) = N/\tau_U + \sum_i^N (V_{i,t} - \mu_U)^2 \quad (43)$$

Which together gives:

$$E_{U,S}((U_t - S_t)^2) + E_U(\sum_i^N (U_t - V_{i,t})^2)$$
$$= (\mu_U - \mu_S)^2 + 1/\tau_U + 1/\tau_S + (N+1)/\tau_U + \sum_i^N (V_{i,t} - \mu_U)^2 \quad (44)$$

#### 2.3 Approximating $q(\theta)$

We will approximate $q_1(\theta)$ with a Normal distribution.

To do this we use a Laplace approximation around the max of $q_1(\theta)$:
$\arg\max(q_1(\theta_t)) = \mu_{\theta,t}$ and with second derivative $-\tau_{\theta,t}$
This gives

$$q_1(\theta_t) \sim \mathcal{N}(\theta_t | \mu_{\theta,t}, 1/\tau_{\theta,t}) \quad (45)$$

##### 2.3.1 First derivative

However in order to find $\arg\max(q_1(\theta_t))$ we differentiate $\log q_1(\theta_t)$ and set equal to 0:

$$d \log q_1(\theta_t)/d\theta_t = -\exp\theta_t E_{U,S}((U_t - S_t)^2)/2$$
$$- \exp\theta_t E_U(\sum_i^N (U_t - V_{i,t})^2)/2$$
$$+ (N+1)/2 - \kappa'(\theta_t - \theta_{t-1}) = 0 \quad (46)$$

$$\exp\theta_t(E_{U,S}((U_t - S_t)^2)/2 + E_U(\sum_i^N (U_t - V_{i,t})^2)/2) = (N+1)/2 - \kappa'(\theta_t - \theta_{t-1}) \quad (47)$$

At this point there is no analytical solution.

##### 2.3.2 Taylor expansion of first derivative

While we could use a numerical approximation for speed of implementation we use Taylor expansion. We need to solve for $\theta$

$$\exp\theta_t(E_{U,S}((U_t-S_t)^2)/2+E_U(\sum_i^N(U_t-V_{i,t})^2)/2)-(N+1)/2+\kappa'(\theta_t-\theta_{t-1})=0 \quad (48)$$

For simplicity we refer to $SV=(E_{U,S}((U_t-S_t)^2)+E_U(\sum_i^N(U_t-V_{i,t})^2))$

We can solve this by using the third order Taylor expansion of the exponential:

$$\begin{aligned}\exp\theta_t &\approx e^{(\theta_*)}+e^{(\theta_*)}(\theta_t-\theta_*)+e^{(\theta_*)}(\theta_t-\theta_*)^2/2+e^{(\theta_*)}(\theta_t-\theta_*)^3/6=\\ &e^{(\theta_*)}\left(1-\theta_*+\theta_*^2/2-\theta_*^3/6+(1-2/2*\theta_*+3/6*\theta_*^2)\theta_t+(1/2-3/6*\theta_*)\theta_t^2+\theta_t^3/6\right)=\\ &e^{(\theta_*)}\left(1-\theta_*+\theta_*^2/2-\theta_*^3/6+(1-\theta_*+\theta_*^2/2)\theta_t+(1/2-\theta_*/2)\theta_t^2+\theta_t^3/6\right) \quad (49)\end{aligned}$$

We can set $\theta_*$ as $\theta_{t-1}$, as we expect that $\theta_t$ will be close to $\theta_*$. For big changes in the variance of $V_i$ this can be off, however this was not a problem in this stimulus set which relied on slow gradual changes.

In order to find $\arg\max(q_1(\theta_t))$ we therefore have to solve

$$\begin{aligned}SV/2{*}e^{(\theta_*)}\left(1-\theta_*+\theta_*^2/2-\theta_*^3/6+(1-\theta_*+\theta_*^2/2)\theta_t+(1/2-\theta_*/2)\theta_t^2+\theta_t^3/6\right)\\ -(N+1)/2+\kappa'(\theta_t-\theta_{t-1})=0 \quad (50)\end{aligned}$$

which can be rewritten as a third order polynomial

$$\theta_t^3+c_1\theta_t^2+c_2\theta_t+c_3=0 \quad (51)$$

where

$$\begin{aligned}c_1&=1/2-\theta_*/2\\ c_2&=1-\theta_*+\theta_*^2+\kappa'/(SV/2*e^{(\theta_*)})\\ c_3&=1-\theta_*+\theta_*^2/2+\theta_*^3/6-(N/2+1/2+\kappa'\theta_{t-1})/(SV/2*e^{(\theta_*)}) \quad (52)\end{aligned}$$

the solution to which, $\theta_t^{optim}$, can be numerically found using Matlab's *nthroot* function.

As we assume that the log-variance only changes slightly between trials the solution closest to the previous value $\theta_{t-1}$ is automatically chosen, $\arg\max(q_1(\theta_t))=\mu_{\theta_t}=\theta_t^{optim}$.

##### 2.3.3 Second derivative

The second derivative is

$$d^2\log q_1(\theta_t)/d\theta_t^2 = -\exp\theta_t * SV/2 - \kappa' \quad (53)$$

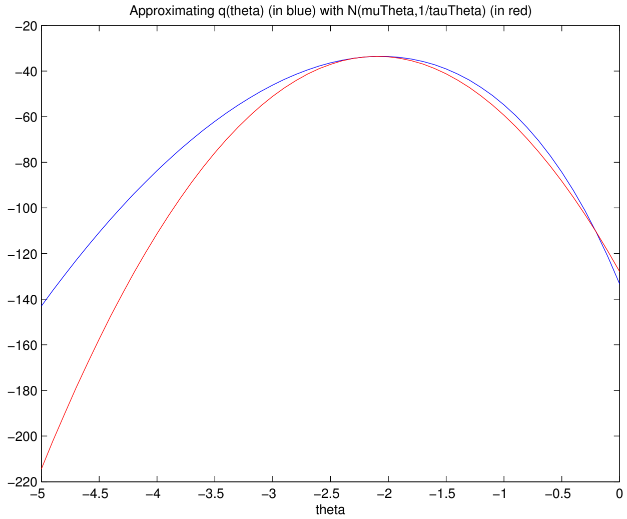

Figure 2: Approximation of theta using Laplace approximation.

We evaluate this at $argmax(q_1(\theta_t))$, so we insert $\theta_t = \mu_{\theta,t}$

Hence we can finally write

$$q_1(\theta_t) \sim \mathcal{N}(\theta_t|\mu_{\theta,t}, 1/\tau_{theta,t}) \tag{54}$$

where

$$\mu_{\theta,t} = \theta_t^{optim} \tag{55}$$

$$\tau_{\theta,t} = \exp(\mu_{\theta,t}) * (SV)/2 + \kappa' \tag{56}$$

and where $SV = (\mu_U - \mu_S)^2 + 1/\tau_S + (N+1)/\tau_U + \sum_i^N (V_{i,t} - \mu_U)^2$

With $q_1(\theta_t)$ a Normal distribution, that makes $q_1(\lambda_{V,t})$ a log-normal distribution with $\mu_{\lambda_{V,t}} = E(\lambda_{V,t}) = E(\exp(\theta_t)) = \exp(\mu_{\theta_t} + 1/(2 * \tau_{\theta_t}))$ (general property of log-normal distribution).

#### 2.4 Final algorithm for C=1

We can now create an iterative algorithm that for each time step $t$ represents the model posterior. Variables $\sigma_A^2 = 1/\lambda_A$, $\sigma_0^2 = 1/\lambda_0$ and $\kappa$ have to be set before hand, together with the input data $A_{1:t}$ and $V_{1:N,1:t}$. For time step $t$:

- 1. initially set

$$\mu_{\theta,t} = \mu_{\theta,t-1}, \tag{57}$$

$$\mu_{S,t} = 0, \tag{58}$$

$$\mu_{U,t} = 1/N \sum V_{i,t}, \tag{59}$$

$$\tau_{\theta,t} = 1 \tag{60}$$

- 2. set $\mu_S, \tau_{S,t}$

$$\mu_{S,t} = (\lambda_A A_t + \exp(\mu_{\theta_t} + 1/(2 * \tau_{\theta_t}))\mu_{U,t})/\tau_{S,t} \tag{61}$$

$$\tau_{S,t} = \lambda_S + \lambda_A + \exp(\mu_{\theta_t} + 1/(2 * \tau_{\theta_t})) \tag{62}$$

- 3. set $\mu_{U,t}, \tau_{U,t}$

$$\mu_{U,t} = (N/(N+1))\bar{V_t} + (1/(N+1))\mu_{S,t} \tag{63}$$

$$\tau_{U,t} = (N+1) * \exp(\mu_{\theta_t} + 1/(2 * \tau_{\theta_t})) \tag{64}$$

where $\bar{V_t} = 1/N \sum_i^N V_{i,t}$

- 4. find $\mu_{\theta_t}$ by solving third order polynomial, equation 51,

$$\mu_{\theta,t} = \theta_{optim} \tag{65}$$

then set $\tau_{\theta_t}$

$$\tau_{\theta,t} = k' + \exp(\mu_{\theta,t}) * ((\mu_{U,t} - \mu_{S,t})^2 + 1/\tau_{S,t} + (N+1)/\tau_{U,t} + \sum_i^N (V_{i,t} - \mu_{U,t})^2)/2 \tag{66}$$

where $\kappa'_t = 1/(1/\kappa + 1/\tau_{\theta,t-1})$

- 5. Repeat steps 2-4 until the change in each parameter is small ($< 0.0001$)

This is then repeated for each time step $t$, providing us with the approximation to the posterior $P_1(S_t, \theta_t, U_t | A_t, V_{1:N,t}) \approx q_1(S_t, U_t, \theta_t)) = q_1(S_t) * q_1(U_t) * q_1(\theta_t)$.

#### 2.5 Posterior for C=2

Due to the independent structure this posterior can be written as

$$P_2(S_{A,t}, S_{V,t}, \lambda_{V,t}, U_t | A_t, V_{1:N,t}) = P(S_{A,t}|A_t) P(S_{V,t}, \lambda_{V,t}, U_t | V_{1:N,t}) \tag{67}$$

where

$$P_2(S_{A,t}|A_t) = P(S_{A,t}) P(A_t|S_{A,t})/Z \tag{68}$$

which is simple enough given the Normal distribution of both $P(S_{A,t})$ and $P(A_t|S_{A,t})$

$$P_2(S_{A,t}|A_t) = \mathcal{N}(S_{A,t}; A_t \sigma_{A0}^2/\sigma_A^2, \sigma_{A0}^2) \tag{69}$$

where $\sigma_{A0}^2 = 1/(1/\sigma_A^2 + 1/\sigma_0^2)$.

Note that for the subject response the posterior $P_2(S_{A,t}|A_t)$ is all that is needed, but for the calculation of the prior $P(\lambda_{V,t})$ for subsequent trial $t+1$ we need to compute the full posterior.

We again use the transformation of parameters

$$\theta_t = \log(\lambda_{V,t}) \tag{70}$$

Proceeding with just the posterior over $S_{V,t}$, $U_t$ and $\theta_t$

$$P_{C=2}(S_{V,t}, \theta_t, U_t|A_{1:t}, V_{1:N,1:t}) \propto P(S_{V,t})P(U_t|S_{V,t}, \theta_t)P(V_{1:N,t}|U_t, \theta_t)$$
$$\int P(\theta_t|\theta_{V,t-1})P(\theta_{V,1:t-1}|A_{1:t-1}, V_{1:N,1:t-1})d\theta_{V,1:t-1}/Z \tag{71}$$

where

$$P(U_t|S_{V,t}, \theta_t) = \mathcal{N}(U_t; S_{V,t}, 1/\exp(\theta_t)) \tag{72}$$

$$P(V_{1:N,t}|U_t, \theta_t) = \prod_n \mathcal{N}(U_t; V_{n,t}, 1/\exp(\theta_t)) \tag{73}$$

$$P(\theta_t|\theta_{t-1}) = \mathcal{N}(\theta_t; \theta_{t-1}, 1/\kappa) \tag{74}$$

We will assume that $P(\theta_{V,1:t-1}|A_{1:t-1}, V_{1:N,1:t-1})$ can be approximated by a Normal distribution (see $q_1(\theta)$ below), thus allowing us to use properties of Normal distributions.

Hence,

$$\int P(\theta_t|\theta_{t-1})P(\theta_{V,1:t-1}|A_{1:t-1}, V_{1:N,1:t-1})d\theta_{V,1:t-1} = \mathcal{N}(\theta_t; \theta_{t-1}, 1/\kappa'_t) \tag{75}$$

where $1/\kappa'_t = 1/\kappa + 1/\tau_{\theta,t-1}$ (due to the convolution of $P(\theta_t|\theta_{t-1})$ with $P(\theta_{t-1}|A_{1:t-1}, V_{1:N,1:t-1})$, both Normal distributed).

While any estimate of $\theta_t$ will depend on $A_{1:t-1}$ and $V_{1:N,1:t-1}$) for ease of notation we will omit those in the following.

The log-posterior is now

$$\log P_2(S_{V,t}, \theta_t, U_t|A_{1:t}, V_{1:N,1:t}) \propto$$
$$-\lambda_0 S_{V,t}^2/2$$
$$-\exp\theta_t(U_t - S_{V,t})^2/2 + \theta_t/2$$
$$-\exp\theta_t \sum_i^N (U_t - V_{i,t})^2/2 + N\theta_t/2$$
$$-\kappa'(\theta_t - \theta_{V,t-1})^2/2 \tag{76}$$

#### 2.6 Variational Bayes approximation for C=2

We will now approximate the log-posterior with variational Bayes by factorization: $P_2(S_t, \theta_t, U_t|A_t, V_{1:N,t}) \approx q_2(S_t, U_t, \theta_t)) = q_2(S_t) * q_2(U_t) * q_2(\theta_t)$ This proceeds similarly to the combined ($C = 1$) model, but with $S_{V,t}$ instead of $S_t$, and with no influence from $A_t$. For completeness the calculations are included here:

#### 2.6.1 $q_2(S_t)$

For $q_2(S_t)$

$$\log q_2(S_{V,t}) \propto -\lambda_0 S_{V,t}^2/2 - E_\theta(\exp(\theta_t))E_U((U_t - S_{V,t})^2)/2 \tag{77}$$

where $E_Y(X)$ signifies the expectation of X over the distribution of Y: $E_Y(X) = \int P(Y)XdY$.

Using $E_U((U_t - V,t)^2) = E_U(U_t^2 + S_{V,t}^2 - 2U_tS_{V,t}) = E_U(U_t^2) + S_{V,t}^2 - 2S_{V,t}E_U(U_t) = (S_{V,t}^2 - S_{V,t}E_U(U_t)*2 + E_U(U_t)^2) - E_U(U_t)^2 + E_U(U_t^2) = (S_{V,t} - E_U(U_t))^2 - E_U(U_t)^2 + E_U(U_t^2)$, where the last two terms do not depend on $S_{V,t}$ (and thus can be discarded) we can rewrite the last term:

$$\log q_2(S_{V,t}) \propto -\lambda_0 S_{V,t}^2/2 - E_\theta(\exp(\theta_t))(S_{V,t} - E_U(U_t))^2/2 \tag{78}$$

#### 2.6.2 $q_2(U_t)$

For $q_2(U_t)$

$$\log q_2(U_t) \propto -E_\theta(\exp(\theta_t))/2\sum_i(U_t - V_{i,t})^2 - E_\theta(\exp(\theta_t))E_S((U_t - S_{V,t})^2)/2 \tag{79}$$

Here we use the same trick

$$\log q_2(U_t) \propto -E_{\theta_t}(\exp(\theta_t))/2\sum_i(U_t - V_{i,t})^2 - E_\theta(\exp(\theta_t))(U_t - E_S(S_{V,t}))^2)/2 \tag{80}$$

###### 2.6.3 $q_2(\theta_t)$

For $q_2(\theta)$

$$\begin{aligned}\log q_2(\theta) = &- \exp\theta_t E_{U,S}((U_t - S_{V,t})^2)/2 \\ &- \exp\theta_t E_U(\sum_i^N(U_t - V_{i,t})^2)/2 \\ &+ (N+1)\log(\exp(\theta))/2 - \kappa'(\theta_t - \theta_{t-1})^2/2\end{aligned} \tag{81}$$

###### 2.6.4 Simplifying $q_2(S_t)$ and $\log q_2(U_t)$

Inspecting $\log q_2(S_{V,t})$ and $\log q_2(U_t)$ we can see that both $q_2(S_{V,t})$ and $q_2(U_t)$ are products of Normal distributions, and thus themselves Normal distributed

$$q_2(S_{V,t}) \sim \mathcal{N}(S_{V,t}|\mu_{S_V,t}, 1/\tau_{S_V,t}) \tag{82}$$

and

$$q_2(U_t) \sim \mathcal{N}(U_t|\mu_{U,t}, 1/\tau_{U,t}) \tag{83}$$

where

$$\mu_{S_V,t} = (\lambda_0 * 0 + E(\exp(\theta_t))\mu_{U,t})/\tau_{S_V,t} \tag{84}$$

$$\tau_{S_V,t} = \lambda_0 + E(\exp(\theta_t)) \quad (85)$$

$$\mu_{U,t} = (E_\theta(\exp(\theta_t))\sum_i^N V_{i,t} + E_\theta(\exp(\theta_t))\mu_{S_V,t})/\tau_{U,t} \quad (86)$$

$$\tau_{U,t} = (N+1) * E_\theta(\exp(\theta_t)) \quad (87)$$

Note that $E_S(S_{V,t}) \approx \int q_2(S_{V,t})S_{V,t}dS_{V,t} = \mu_{S_V,t}$ and $E_U(U_t) \approx \int q_2(U_t)U_t dU_t = \mu_{U,t}$

We can approximate $q_2(\theta)$ with a Normal distribution, in exactly the same way as for C=1. As equations are identical (see above) they will not be repeated here.

#### 2.7 Final algorithm for C=2

We can now create an iterative algorithm that for each time step $t$ represents the variational Bayes approximation of the model posterior over $S_{V,t}, U_t$ and $\lambda_{V,t}$ (or rather $\theta_t$):, $P(S_{V,t}, U_t, \theta_t | V_{1:N,t})$. Variable $\kappa$ has to be set before hand, together with the input data $A_{1:t}$ and $V_{1:N,1:t}$. For time step $t$:

- 1. initially set $\mu_{\theta,t} = \mu_{\theta,t-1}$, $\mu_{U,t} = 1/N\sum V_{i,t}$, and $\tau_{\theta,t} = 1$
- 2. set $\mu_{S_V}, \tau_{S_V,t}$

$$\mu_{S_V,t} = (\exp(\mu_{\theta_t} + 1/(2 * \tau_{\theta,t}))\mu_{U,t})/\tau_{S_V,t} \quad (88)$$

$$\tau_{S_V,t} = \lambda_0 + \exp(\mu_{\theta_t} + 1/(2 * \tau_{\theta,t})) \quad (89)$$

- 3. set $\mu_U, \tau_{U,t}$

$$\mu_{U,t} = \Big(N\bar{V}_t + \mu_{S_V,t}\Big)/(N+1) \quad (90)$$

$$\tau_{U,t} = (N+1) * \exp(\mu_{\theta_t} + 1/(2 * \tau_{\theta,t})) \quad (91)$$

  where $\bar{V}_t = 1/N\sum_i^N V_{i,t}$

- 4. find $\mu_{\theta,t}$ by numerically solving polynomial, equation 51,

$$\mu_{\theta,t} = \theta_{optim} \quad (92)$$

  then set $\tau_{\theta,t}$

$$\tau_{\theta,t} = k' + \exp(\mu_{\theta,t}) * \Big((\mu_{U,t} - \mu_{S_V,t})^2 + 1/\tau_{S_V,t} + (N+1)/\tau_{U,t} + \sum_i^N (V_{i,t} - \mu_{U,t})^2\Big)/2 \quad (93)$$

  where $\kappa'_t = 1/(1/\kappa + 1/\tau_{\theta,t-1})$

- 5. Repeat steps 2-4 until convergence, i.e. until the change in each parameter is small ($< 0.0001$)

This is then repeated for each time step $t$, providing us with the approximation to the posterior $P_2(S_t, \theta_t, U_t | A_t, V_{1:N,t}) \approx q_2(S_t, U_t, \theta_t)) = q_2(S_t) * q_2(U_t) * q_2(\theta_t)$.

See Fig. 3 below for an example of the learned inference of the visual variance $\sigma^2_{V,t} = 1/\lambda_{V,t} \sim 1/\log q_2(\theta_t)$, compared with a simple instantaneous learner model that assumes $\sigma^2_{V,t} = 1/N \sum_i (V_{i,t} - \bar{V}_t)^2$.

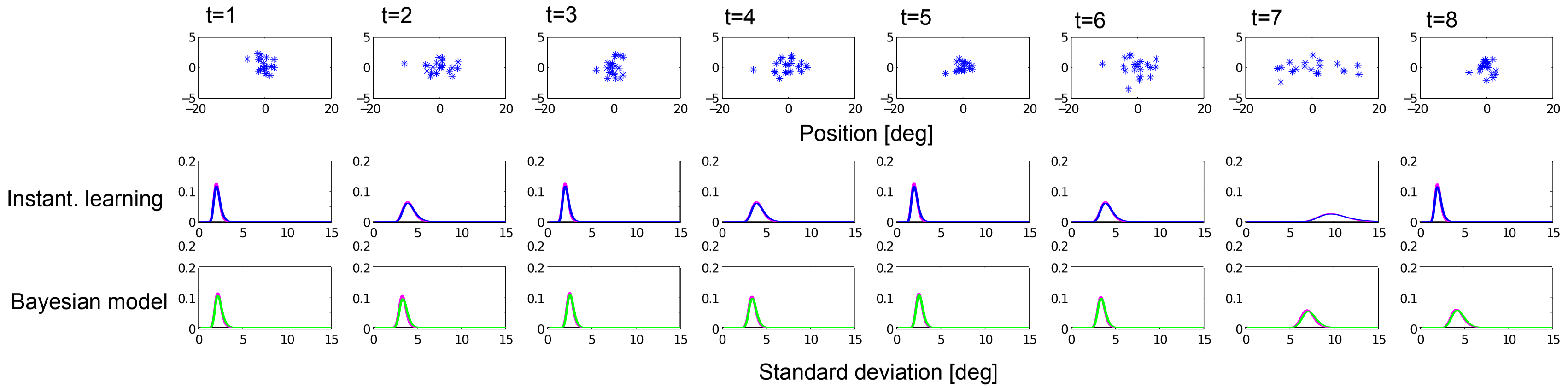

Figure 3: Compring varitional Bayes approximation with a numerical discretised grid approximation. Top row: Example visual stimuli over eight subsequent trials. Middle row: The distribution of estimated sample variance, with no learning over trials. Bottom row: The distribution of $\sigma_{V,t}$ for the Bayesian model that incorporates the learning across trials. Red line is the numerical comparison when using a discretised grid to estimate variance, as opposed to the variational Bayes (green line).

### 3 Marginal model evidence

Recall that the posterior is a mixture of the forced fusion and independent solutions, with the mixture determined by the posterior probability of either model generating the data:

$$P(C_t = 1|A_t, V_{1:N,t}) = \frac{P(A_t, V_{1:N,t}|C_t = 1)P(C_t = 1)}{P(A_t, V_{1:N,t}|C_t = 1)P(C_t = 1) + P(A_t, V_{1:N,t}|C_t = 2)P(C_t = 2)} \tag{94}$$

To evaluate this we need to calculate the marginal model evidence, $P(A_t, V_{1:N,t}|C_t)$, for either model.

One way to do so is by a sampling approximation, but here we utilise the variational results we have already found.

#### 3.1 Model likelihood for C=2, two sources $S_{V,t}, S_{A,t}$

We need to evaluate the model likelihood for both $C = 1$ and $C = 2$. The case for $C = 2$ is slightly simpler, hence we start with this:

$$\begin{aligned} P(A_t, V_{1:N,t}|C_t = 2) = P(A_t|C_t)P(V_{1:N,t}|C_t = 2) = \\ \int P(A_t|S_{A,t}, C_t = 2)P(S_{A,t}|C_t = 2)dS_{A,t}* \\ \int P(V_{1:N,t}|U_t, \lambda_t, C_t = 2)P(U_t, \lambda_t, S_{V,t}|C_t = 2)dVd\lambda dS_{V,t} \end{aligned} \tag{95}$$

The first integral is easy as it is just the integral of the product of two Normal distributions.

$$\int P_2(A_t|S_{A,t})P_2(S_{A,t})dS = \frac{1}{\sqrt{(2\pi(\sigma_A^2 + 1/\tau_0)}} \exp(-\frac{(A - \mu_0)^2}{2(\sigma_A^2 + 1/\tau_0)}) \tag{96}$$

It is however more convenient to operate in log-space

$$\log P_2(A_t) = -\log(\sqrt{2\pi(\sigma_A^2 + 1/\tau_0)}) - (A - \mu_0)^2/(2(\sigma_A^2 + 1/\tau_0)) \tag{97}$$

The second integral we approximate through the Free Energy that we already maximise iteratively in the variational Bayes algorithm.

$$\begin{aligned} \log P(V_{1:N,t}|C_t = 2) = \log \int P(V_{1:N,t}|U_t, \theta_t, S_{V,t}, C_t = 2)P(U_t, \theta_t, S_{V,t}|C_t = 2)dU_t d\theta_t dS_{V,t} \\ \approx L_2(q) = \int q_2(U_t, \theta_t, S_{V,t})* \\ \log \frac{P_2(V_{1:N,t}|U_t, \theta_t, C_t)P_2(U_t|, S_{V,t}, \theta_t, C_t)P_2(S_{V,t}|C_t)P_2(\theta_t|C_t)}{q_2(U_t, \theta_t, S_{V,t})} dU_t d\theta_t dS_{V,t} \end{aligned} \tag{98}$$

(this approximation becomes exact if the variational approximation is exact, ie if the Kulback-Leibler difference between the posterior $P_2(U_t, \theta_t, S_{V,t}|V_{1:N})$ and the approximation $q_2(U_t, \theta_t, S_{V,t})$ becomes zero.)

This can be interpreted as taking the expectation with regards to the posterior approximation, and due to the properties of the logarithm this can be separated into a sum of expectations:

$$\begin{aligned} L_2(q) = E(\log P_2(V_{1:N,t}|U_t, \theta_t)) + \\ E(\log P_2(U_t|S_{V,t}, \theta_t) + E(\log P_2(S_{V,t})) + E(\log P_2(\theta_t)) \\ - E(\log q_2(U_t)) - E(\log q_2(\theta_t)) - E(\log q_2(S_{V,t})) \end{aligned} \quad (99)$$

where (due to eq. 43)

$$\begin{aligned} E(\log P_2(V_{1:N,t}|U_t, \theta_t)) = E(\log \prod_i P_2(V_{i,t}|U_t, \theta)) = \\ E(\log \prod_i \sqrt{\frac{\exp(\theta_t)}{2\pi}} \exp(-(V_{i,t} - U_t)^2 \exp(\theta_t)/2) = \\ E(\sum_i \log(\sqrt{\frac{\exp(\theta_t)}{2\pi}}) - (V_{i,t} - U_t)^2 \exp(\theta_t)/2) = \\ N/2(E_{\theta_t}(\theta) - \log(2\pi)) + E_{U_t}(\sum_i -(V_{i,t} - U_t)^2))E(\exp(\theta_t))/2 = \\ N/2(\mu_\theta - \log(2\pi)) - (N/\tau_{U,t} + \sum_i (V_{i,t} - \mu_U)^2) \exp(\mu_{\theta,t} + 1/(2\tau_{\theta,t}))/2 \end{aligned} \quad (100)$$

and (since $E(X^2) = \mu_X^2 + \sigma_X^2$)

$$\begin{aligned} E(\log P_2(U_t)|S_{V,t}, \theta_t)) = E(\log \mathcal{N}(U_t; S_{V,t}, 1/\exp(\theta_t))) = \\ E(\log \sqrt{\frac{\exp \theta_t}{2\pi}} \exp(-(U_t - S_{V,t})^2) \exp(\theta_t)/2)) = \\ (\mu_{\theta,t} - \log(2\pi))/2 - [(\mu_U - \mu_{S_V,t})^2 + 1/\tau_{U,t} + 1/\tau_{S_V,t}] \exp(\mu_{\theta,t} + 1/(2\tau_{\theta,t}))/2 \end{aligned} \quad (101)$$

and

$$\begin{aligned} E(\log P_2(S_{V,t})) = E(\log \mathcal{N}(S_{V,t}; \mu_0, \sigma_0^2)) = \\ E(\log \sqrt{\frac{1}{\sigma_0^2 2\pi}} \exp(-(S_{V,t}-\mu_0)^2)/(2\sigma_0^2)) = -\log(\sigma_0^2 2\pi)/2 - [(\mu_{S_V,t}-\mu_0)^2 + 1/\tau_{S_V,t}]/(2\sigma_0^2) \end{aligned} \quad (102)$$

and (due to eq. 75)

$$E(\log P_2(\theta_t)) = E(\ \mathcal{N}(\theta_t;\theta_{t-1},1/\kappa')) =$$

$$E(\log\sqrt{\frac{\kappa'}{2\pi}}\exp(-(\theta_t-\mu_{\theta,t-1})^2\kappa'/2)) = (\log\kappa'-\log(2\pi))/2-[(\mu_{\theta,t}-\mu_{\theta,t-1})^2+1/\tau_{\theta,t}]\kappa'/2 \tag{103}$$

and

$$\begin{aligned} E(\log q_2(U_t)) = E(\log\ \mathcal{N}(U_t;\mu_{U,t},1/\tau_{U,t})) = \\ E(\log(\sqrt{\frac{\tau_{U,t}}{2\pi}}\exp(-(U_t-\mu_{U,t})^2\tau_{U,t}/2))) = \\ \log\sqrt{\frac{\tau_{U,t}}{2\pi}} - E((U_t-\mu_{U,t})^2\tau_{U,t}/2) = \\ (\log\tau_{U,t}-\log(2\pi))/2-(E(U_t^2)+\mu_{U,t}^2-2\mu_{U,t}E(U_t))\tau_{U,t}/2 = \\ (\log\tau_{U,t}-\log(2\pi))/2-(\mu_{U,t}^2+1/\tau_{U,t}^2+\mu_{U,t}^2-2\mu_{U,t}\mu_{U,t})\tau_{U,t}/2 = \\ (\log\tau_{U,t}-\log(2\pi)-1)/2 \end{aligned} \tag{104}$$

and

$$\begin{aligned} E(\log q_2(S_t)) = E(\log\ \mathcal{N}(S_{V,t};\mu_{S_V,t},1/\tau_{S_V,t})) = \\ E(\log\sqrt{\frac{1}{2\pi\sigma_0^2}}\exp(-(S_{V,t}-\mu_{S_V,t})^2\tau_{S_V,t}/2)) = \\ (\log\tau_{S_V,t}-\log(2\pi)-1)/2 \end{aligned} \tag{105}$$

and

$$\begin{aligned} E(\log q_2(\theta_t)) = E(\log\ \mathcal{N}(\theta_t;\mu_{\theta,t},1/\tau_{\theta,t})) = \\ (\log\tau_{\theta,t}-\log(2\pi)-1)/2 \end{aligned} \tag{106}$$

In total we now have

$$
\begin{aligned}
\log P_2(A_t, V_{1:N,t}|C_t) \approx \log P_2(A_t) + L_2 = \\
-(\log(2\pi) + \log(\sigma_A^2 + 1/\tau_0))/2 - (A - \mu_0)^2/(2(\sigma_A^2 + 1/\tau_0)) \\
+ (\mu_{\theta,t} - \log(2\pi))N/2 - [N/\tau_{U,t} + \sum_i^N (V_{i,t} - \mu_U)^2] \exp(\mu_{\theta,t} + 1/(2\tau_{\theta,t}))/2 \\
+ ((\mu_{\theta,t} - \log(2\pi))/2 - [(\mu_U - \mu_{S_V,t})^2 + 1/\tau_{U,t} + 1/\tau_{S_V,t}] \exp(\mu_{\theta,t} + 1/(2\tau_{\theta,t}))/2 \\
- \log(\sigma_0^2 2\pi)/2 - [(\mu_{S_V,t} - \mu_0)^2 + 1/\tau_{S_V,t}]/(2\sigma_0^2) \\
+ (\log\kappa' - \log(2\pi))/2 - [(\mu_{\theta,t} - \mu_{\theta,t-1})^2 + 1/\tau_{\theta,t}]\kappa'/2 \\
- (\log\tau_{U,t} - \log(2\pi) - 1)/2 \\
- (\log\tau_{S_V,t} - \log(2\pi) - 1)/2 \\
- (\log\tau_{\theta,t} - \log(2\pi) - 1)/2
\end{aligned}
\tag{107}
$$

Although lengthy, this is trivial and fast to compute numerically in Matlab (e.g.). Note that all estimates come from the variational Bayes approximation $q_2(S_t, U_t, \theta_t)$.

#### 3.2 Model likelihood for C=1, one source $S_t = S_{V,t} = S_{A,t}$

We now need to do the same for the one source model.

$$
\begin{aligned}
P(A_t, V_{1:N,t}|C_t = 1) = P_1(A_t, V_{1:N,t}) = \\
\int P_1(A_t|S_t, C_t = 1) P_1(V_{1:N,t}|U_t, \lambda_t, C_t = 1) P_1(U_t, \lambda_t, S_t|C_t = 1) dV d\lambda dS_t
\end{aligned}
\tag{108}
$$

Note that for simplicity in notation we will use $P_1$ to indicate the probability within the model given $C_t = 1$

We again approximate through the Free Energy that we already maximised iteratively in the variational Bayes algorithm.

$$
\begin{aligned}
\log P_1(A_t, V_{1:N,t}) = \log \int P_1(V_{1:N,t}|U_t, \theta_t, S_t) P_1(A_t|S_t) P_1(U_t, \theta_t, S_t) dU_t d\theta_t dS_t \\
\approx L_{C_t=1}(q_1) = \\
\int q_1(U_t, \theta_t, S_{V,t}) \log \frac{P_1(V_{1:N,t}|U_t, \theta_t) P_1(A_t|S_t) P_1(U_t|, S_t, \theta_t) P_1(S_t) P_1(\theta_t)}{q_1(U_t, \theta_t, S_t)} dU_t d\theta_t dS_t
\end{aligned}
\tag{109}
$$

(this approximation becomes exact if the variational approximation is exact, ie if the Kulback-Leibler difference between the posterior $P_1(V, \theta_t, S_{V,t}|V_{1:N,t})$ and the approximation $q_1(U_t, \theta_t, S_{V,t})$ becomes zero.)

This can be interpreted as taking the expectation with regards to the posterior approximation, and due to the properties of the logarithm this can be separated into a sum of expectations:

$$
\begin{aligned}
L_1(q) = E(\log P_1(A_t|S_t)) + E(\log P_1(V_{1:N,t}|U_t, \lambda_t)) \\
+ E(\log P_1(U_t)) + E(\log P_1(\theta_t)) + E(\log P_1(S_{V,t})) \\
- E(\log q_1(U_t)) - E(\log q_1(\theta_t)) - E(\log q_1(S_{V,t}))
\end{aligned} \quad (110)
$$

where (since $E(X^2) = \mu_X^2 + \sigma_X^2$)

$$
\begin{aligned}
&E(\log P_1(A_t|S_t)) = \\
&\quad E(\log \sqrt{\frac{1}{2\pi\sigma_A^2}} \exp(-(A_t - S_t)^2/(2\sigma_A^2)) = \\
&\quad - \log(\sigma_A^2 2\pi)/2 - [(A_t - \mu_S)^2 + 1/\tau_{S,t}]/(2\sigma_A^2)
\end{aligned} \quad (111)
$$

$$
\begin{aligned}
&E(\log P_1(V_{1:N,t}|U_t, \theta_t)) = E(\log \prod_i P_1(V_{i,t}|U_t, \theta)) = \\
&E(\log \prod_i \sqrt{\frac{\exp(\theta_t)}{2\pi}} \exp(-(V_{i,t} - U_t)^2 \exp(\theta_t)/2) = \\
&E(\sum_i \log(\sqrt{\frac{\exp(\theta_t)}{2\pi}}) - (V_{i,t} - U_t)^2 \exp(\theta_t)/2) = \\
&n/2(E_{\theta_t}(\theta) - \log(2\pi)) + E(\exp(\theta_t))/2E_{U_t}(\sum_i -(V_{i,t} - U_t)^2)) = \\
&n/2(\mu_{\theta,t} - \log(2\pi)) + \exp(\mu_{\theta,t} - 1/(2\tau_{\theta,t}))/2(N/\tau_{U,t} + \sum_i (V_{i,t} - \mu_U)^2)
\end{aligned} \quad (112)
$$

and

$$
\begin{aligned}
&E(\log P_1(U_t)|S_t, \theta_t)) = E(\log \ \mathcal{N}(U_t; S_t, 1/exp(\theta_t))) = \\
&E(\log \sqrt{\frac{\exp \theta_t}{2\pi}} \exp(-(U_t - S_t)^2) \exp(\theta_t)/2)) = \\
&(\mu_{\theta,t} - \log(2\pi))/2 - [(\mu_U - \mu_{S,t})^2 + 1/\tau_{U,t} + 1/\tau_{S,t}] \exp(\mu_{\theta,t} + 1/(2\tau_{\theta,t}))/2
\end{aligned} \quad (113)
$$

and

$$
\begin{aligned}
&E(\log P_1(S_t)) = E(\log \ \mathcal{N}(S_t; \mu_0, \sigma_0^2)) = \\
&E(\log \sqrt{\frac{1}{\sigma_0^2 2\pi}} \exp(-(S_t - \mu_0)^2)/(2\sigma_0^2)) = -\log(\sigma_0^2 2\pi)/2 - [(\mu_{S,t} - \mu_0)^2 + 1/\tau_{S,t}]/(2\sigma_0^2)
\end{aligned} \quad (114)
$$

and (due to eq. 75)

$$E(\log P_1(\theta_t)) = E(\ \mathcal{N}(\theta_t;\theta_{t-1},1/\kappa')) =$$
$$E(\log\sqrt{\frac{\kappa'}{2\pi}}\exp(-(\theta_t-\mu_{\theta,t-1})^2\kappa'/2)) = (\log\kappa'-\log(2\pi))/2-[(\mu_{\theta,t}-\mu_{\theta,t-1})^2+1/\tau_{\theta,t}]\kappa'/2 \tag{115}$$

and

$$\begin{aligned}E(\log q_1(U_t)) = E(\log\ \mathcal{N}(U_t;\mu_{U,t},1/\tau_{U,t})) =\\ E(\log\sqrt{\frac{\tau_{U,t}}{2\pi}}\exp(-(U_t-\mu_{U,t})^2\tau_{U,t}/2)) =\\ \log\sqrt{\frac{\tau_{U,t}}{2\pi}} - E((U_t-\mu_{U,t})^2\tau_{U,t}/2) =\\ (\log\tau_{U,t}-\log(2\pi))/2-(E(U_t^2)+\mu_{U,t}^2-2\mu_{U,t}E(U_t))\tau_{U,t}/2 =\\ (\log\tau_{U,t}-\log(2\pi)-1)/2\end{aligned} \tag{116}$$

and

$$\begin{aligned}E(\log q_1(S_t)) = E(\log\ \mathcal{N}(S_t;\mu_{S,t},1/\tau_{S,t})) =\\ E(\log\sqrt{\frac{1}{2\pi\sigma_0^2}}\exp(-(S_t-\mu_{S,t})^2\tau_{S,t}/2)) =\\ (\log\tau_{S,t}-\log(2\pi)-1)/2\end{aligned} \tag{117}$$

and

$$\begin{aligned}E(\log q_1(\theta_t)) = E(\log\ \mathcal{N}(\theta_t;\mu_{\theta,t},1/\tau_{\theta,t})) =\\ (\log\tau_{\theta,t}-\log(2\pi)-1)/2\end{aligned} \tag{118}$$

In total we now have

$$\begin{aligned}\log P_1(A_t,V_{1:N,t}) \approx\\ L_1(q_1) = -\log(\sigma_A^2 2\pi)/2-[(A_t-\mu_S)^2+1/\tau_{S,t}]/(2\sigma_A^2)\\ +N/2(\mu_{\theta,t}-\log(2\pi))-(N/\tau_{U,t}+\sum_i^N(V_{i,t}-\mu_U)^2)\exp(\mu_{\theta,t}+1/(2\tau_{\theta,t}))/2\\ +((\mu_{\theta,t}-\log(2\pi))/2-[(\mu_U-\mu_{S_V,t})^2+1/\tau_{U,t}+1/\tau_{S_V,t}]\exp(\mu_{\theta,t}+1/(2\tau_{\theta,t}))/2\\ -\log(\sigma_0^2 2\pi)/2-[(\mu_{S_V,t}-\mu_0)^2+1/\tau_{S_V,t}]/(2\sigma_0^2)\\ +(\log\kappa'-\log(2\pi))/2-[(\mu_{\theta,t}-\mu_{\theta,t-1})^2+1/\tau_{\theta,t}]\kappa'/2\\ -(\log\tau_{U,t}-\log(2\pi)-1)/2\\ -(\log\tau_{S,t}-\log(2\pi)-1)/2\\ -(\log\tau_{\theta,t}-\log(2\pi)-1)/2\end{aligned} \tag{119}$$

(where all first and second order moments $(\mu, \tau)$ have been derived from $q_1$).

This is identical to the result from $C = 2$ except for the first line.

In total this provides us with an approximation to the model evidence for each model, $P(A_t, V_{1:N,t}|C_t = 1)$ and $P(A_t, V_{1:N,t}|C_t = 2)$, which can be used to calculate the posterior probability of either model given data, $P(C_t|A_{1:t}, V_{1:N,1:t})$.

### 4 Putting it all together

For either sub-model the factorization (due to assumptions and variational Bayes approximation) allows us to write out the equations for the variable for subject choice:

$$
\begin{aligned}
&P(S_t|A_{1:t}, V_{1:N,1:t}) = \\
&\quad P(S_t|A_{1:t}, V_{1:N,1:t}, C_t = 1)P(C_t = 1|A_{1:t}, V_{1:N,1:t}) + \\
&\quad P(S_{A,t}|A_{1:t}, V_{1:N,1:t}, C_t = 2)P(C_t = 2|A_{1:t}, V_{1:N,1:t}) \approx \\
&\quad q_{C=1,t}(S_t)P(C_t = 1|A_{1:t}, V_{1:N,1:t}) + \\
&\quad P(S_{A,t}|A_{1:t}, C_t = 2)P(C_t = 2|A_{1:t}, V_{1:N,1:t})
\end{aligned} \tag{120}
$$

This is now a mixture of two Gaussian distributions (due to the Variational Bayes approximation), with mixture weights given by the model evidence (partly approximated by the Free Energy).

We will assume subjects report the mean of the distribution i.e.

$$
\hat{S}_t = \hat{S}_{C=1,t}P(C_t = 1|A_{1:t}, V_{1:N,1:t}) + \hat{S}_{C=2,t}P(C_t = 2|A_{1:t}, V_{1:N,1:t}) \tag{121}
$$

where $\hat{S}_{C=1,t} = \mu_{S,t}$ for $C = 1$ and $\hat{S}_{C=2,t} = \mu_{S,A,t}$ for $C = 2$.

We also need a prior over the visual log-reliability for the following trial

$$
\begin{aligned}
&P(\theta_t|A_{1:t}, V_{1:N,1:t}) = \\
&\quad P(\theta_t|A_{1:t}, V_{1:N,1:t}, C = 1)P(C = 1|A_{1:t}, V_{1:N,1:t}) + \\
&\quad P(\theta_t|A_{1:t}, V_{1:N,1:t}, C = 2)P(C = 2|A_{1:t}, V_{1:N,1:t})
\end{aligned} \tag{122}
$$

While this is a mixture of two Gaussians, we need the prior to be a single Gaussian in order for our approximation scheme above to work. We will approximate this mixture with a single Gaussian (essentially fitting a Gaussian to the mixture of two Gaussians).

$$
P(\theta_t|A_{1:t}, V_{1:N,1:t}) = \mathcal{N}(\theta_t|\mu_{\theta,t}, 1/\tau_{\theta,t}) \tag{123}
$$

where (due to the first and second order moments of mixture distributions)

$$
\begin{aligned}
\mu_{\theta,t} = \mu_{\theta,t,C=1}&P(C_t = 1|A_{1:t}, V_{1:N,1:t}) + \\
&\mu_{\theta,t,C=2}P(C_t = 2|A_{1:t}, V_{1:N,1:t})
\end{aligned} \tag{124}
$$

$$
\begin{aligned}
1/\tau_{\theta,t} = (1/\tau_{\theta,t,C=1} + \mu^2_{\theta,t,C=1})P(C_t = 1|A_{1:t}, V_{1:N,1:t}) \\
+ (1/\tau_{\theta,t,C=2} + \mu^2_{\theta,t,C=2})P(C_t = 2|A_{1:t}, V_{1:N,1:t}) \\
- \mu^2_{\theta,t} \quad (125)
\end{aligned}
$$

While fitting a Gaussian to the sum of two Gaussians could be a very in-exact approximation, in practice the two individual distributions are close enough for this not to be a problem (as any contribution from $A_t$ to the posterior of $\theta_t$ is very small).

In **conclusion**, subjects report $\hat{S}_t$ (through a button response, see eq. 121) and they propagate the posterior $P(\theta_t|A_{1:t}, V_{1:N,1:t})$ (see eq. 123) as prior for the next trial.
